# IRF3 regulates neuroinflammatory responses and the expression of genes associated with Alzheimer’s disease

**DOI:** 10.1101/2024.03.08.582968

**Authors:** Radhika Joshi, Veronika Brezani, Gabrielle M Mey, Sergi Guixé-Muntet, Marti Ortega-Ribera, Yuan Zhuang, Adam Zivny, Sebastian Werneburg, Jordi Gracia-Sancho, Gyongyi Szabo

## Abstract

The pathological role of interferon signaling is emerging in neuroinflammatory disorders, yet, the specific role of Interferon Regulatory Factor 3 (IRF3) in neuroinflammation remains poorly understood. Here, we show that global IRF3 deficiency delays TLR4-mediated signaling in microglia and attenuates the hallmark features of LPS-induced inflammation such as cytokine release, microglial reactivity, astrocyte activation, myeloid cell infiltration, and inflammasome activation. Moreover, expression of a constitutively active IRF3 (S388D/S390D:IRF3-2D) in microglia induces a transcriptional program reminiscent of the Activated Response Microglia and the expression of genes associated with Alzheimer’s Disease, notably *apolipoprotein-e*. Lastly, using bulk-RNAseq of IRF3-2D brain myeloid cells, we identified Z-DNA binding protein-1 as a target of IRF3 that is relevant across various neuroinflammatory disorders. Together, our results identify IRF3 as an important regulator of LPS-mediated neuroinflammatory responses and highlight IRF3 as a central regulator of disease-specific gene activation in different neuroinflammatory diseases.

## Introduction

Type I interferon (IFN-I) signaling is a critical adaptive immune response best known to combat viral infections ^1, 2^. The role of IFN-I signaling in the regulation of innate immunity and sterile inflammatory conditions is increasingly recognized. The pathological role of interferon signaling has been reported in a variety of neurological disorders including Alzheimer’s disease (AD), Down syndrome, traumatic brain injury (TBI), and stroke ^3–8^. Interferon signaling is also associated with behavioral changes such as cognitive decline, anxiety, depression, and susceptibility to stress ^9–11^. Interferonopathies are another class of neuropathological disorders specifically classified as such based on their excessive activation of interferon signaling ^12^. Relevant to the role of IFN-I, single nucleotide polymorphisms in interferon-stimulated genes (ISGs) have been associated with AD ^13^.

Single-cell RNA sequencing techniques have discovered interferon-responsive microglia (IRMs with antiviral immune response) in diseases such as AD, multiple sclerosis and during natural aging ^14–16^. IFN-responsive astrocytes and oligodendrocytes have also been described in AD models and aging ^17, 18^. However, a comprehensive understanding of the underlying molecular mechanisms and function of these cell types is still under investigation.

Interferon signaling is regulated via 9 transcription factors called interferon response factors IRF1-9 ^19^. Among these, IRF3 is at the crossroads of adaptive and innate immune responses. IRF3 activation is triggered downstream of TLR3, RIG-I, and MDA-5 in response to dsRNA, typically observed during viral infections ^19^. IRF3 is also activated downstream of TLR4 in a MyD88 independent fashion involving the TRIF adapter molecule ^20, 21^. Following TLR3/4 activation, IRF3 undergoes phosphorylation and dimerization leading to nuclear entry that drives the expression of ISGs ^19, 21^. While IRF3-mediated signaling has been well-studied in various models of peripheral inflammation ^20, 22–25^, in-depth studies directly investigating the role of IRF3 in neuroinflammatory conditions are lacking.

In this study, we examined the direct consequences of IRF3 perturbations on neuroinflammation and microglia. We used the commonly used model of neuroinflammation induced by lipopolysaccharides (LPS), to mimic TLR4 activation. We observed that IRF3 plays a critical role in various features of LPS-mediated proinflammatory changes such as sickness behavior, cytokine production, myeloid cell infiltration and inflammasome activation. Furthermore, we showed that the mere expression of a constitutively active form of IRF3 (IRF3-2D) is sufficient to trigger a proinflammatory phenotype in microglia reminiscent of the IRMs. Importantly, IRF3 activation leads to the expression of genes associated with AD, most notably, apolipoprotein-e (*apoe)*. Lastly, we compared the transcriptome of brain myeloid cells from the IRF3-2D mouse model to that of other neuroinflammatory conditions. We identified Zbp1 as one of the common proinflammatory signatures in microglia across different neurological disorders and we show that IRF3 directly regulates Zbp1. Taken together, we demonstrate that IRF3 plays an important role in proinflammatory responses induced by LPS. Furthermore, selective activation of IRF3 induces features of IRM and certain AD-associated genes.

## Results

### IRF3KO mice show attenuated sickness behavior and reduced proinflammatory and IFN responses after an acute LPS challenge

To determine the relative contribution of the IRF3-induced signaling cascade on the proinflammatory effects of LPS, we first administered LPS (1mg/kg) to wild type (WT) and IRF3KO (whole body knockout) mice and euthanized mice 6 hours later. We analyzed LPS-induced sickness behavior (Fig 1A) in the open field test and observed that WT mice showed reduced locomotion and velocity ∼5h after LPS administration compared to the vehicle group. This reduction in activity was significantly attenuated in the IRF3KO-LPS treated group (Fig 1B). Since sickness behavior is correlated to the peripheral and central nervous system (CNS) release of cytokines such as IL1β, TNFα, IL6 (Salvador et al 2021, Dantzer et al 2009), we assessed the levels of different cytokines and chemokines in cortical lysates to assess the state of neuroinflammation. We found a significant upregulation in IL1β, IL6, IL1α, MCP1, and CXCL1 levels in the cortices of mice treated with LPS in the WT group (Fig 1C). However, cytokine (IL1β, IL1α, IL6) and chemokine (MCP1, CXCL1) induction by LPS was absent or significantly attenuated in IRF3KO mice (Fig 1C).

**Figure 1:**
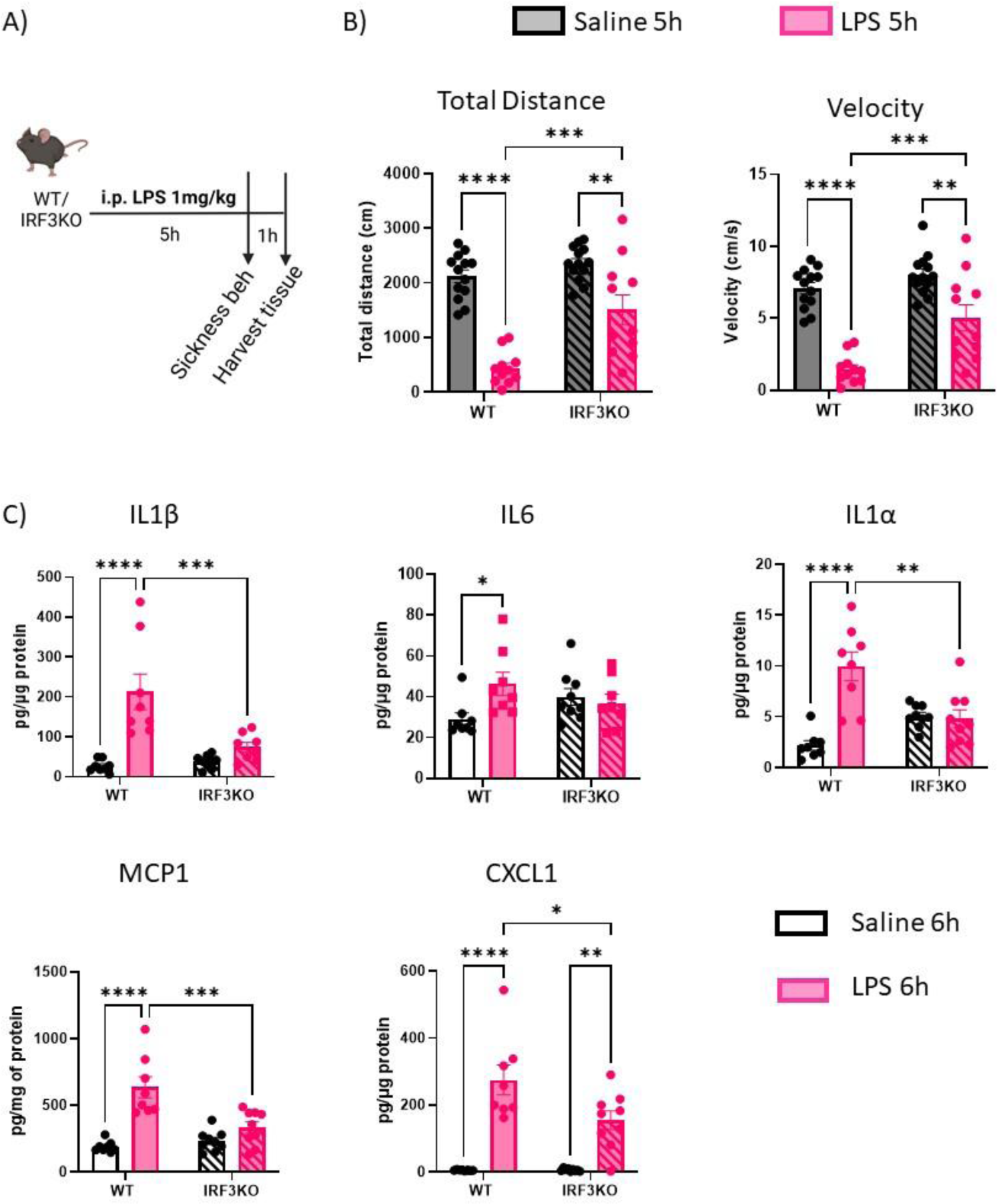
IRF3 deletion attenuates the proinflammatory effects of acute LPS challenge. A) Schematic of the acute LPS challenge model. Sickness behavior was recorded in the open field arena ∼5h after i.p. (intraperitoneal) LPS administration and tissue was collected after 6h. B) Quantification of distance traveled and velocity of movement in open field arena shows that IRF3KO mice display attenuated sickness behavior compared to the WT. N=11-13 for each group. C) Quantification of Elisa from cortical lysates shows that proinflammatory cytokines are significantly upregulated in WT mice on LPS challenge but remain significantly reduced in the IRF3KO cortices compared to the WT. N=11-13 for each group Two-way ANOVA with Tukey’s multiple comparisons. *p<0.05, **p<0.01, ***p<0.001, ****p<0.0001

Since microglia are the key mediators of proinflammatory responses, we tested the expression of proinflammatory transcripts in flow-sorted microglia (CD11b^+^,CD45^intermediate^)(Fig 2A). IRF3 is critical for interferon responses downstream of TLR4, thus we first assessed signatures of interferon signaling followed by other proinflammatory mediators. LPS stimulation induced signatures of interferon signaling (*Ifit1*, *Isg15, Gbp2*) in WT mice, but ISG expression after LPS treatment was abrogated in IRF3KO mice (Fig 2B-D). Similarly, proinflammatory transcripts of *Cox2* and *H2-D1* were significantly increased in microglia of the WT-LPS but not in the IRF3KO-LPS group (Fig 2E,F). Interestingly, IRF3KO mice showed more sensitivity to LPS-induced *C3* transcripts compared to the WT (Fig 2G) and downregulation of the homeostatic marker *P2ry12* was comparable between WT and IRF3KO after acute LPS injection (Fig H). These data collectively suggested that IRF3 partially contributes to the proinflammatory effects of LPS in microglia.

**Figure 2:**
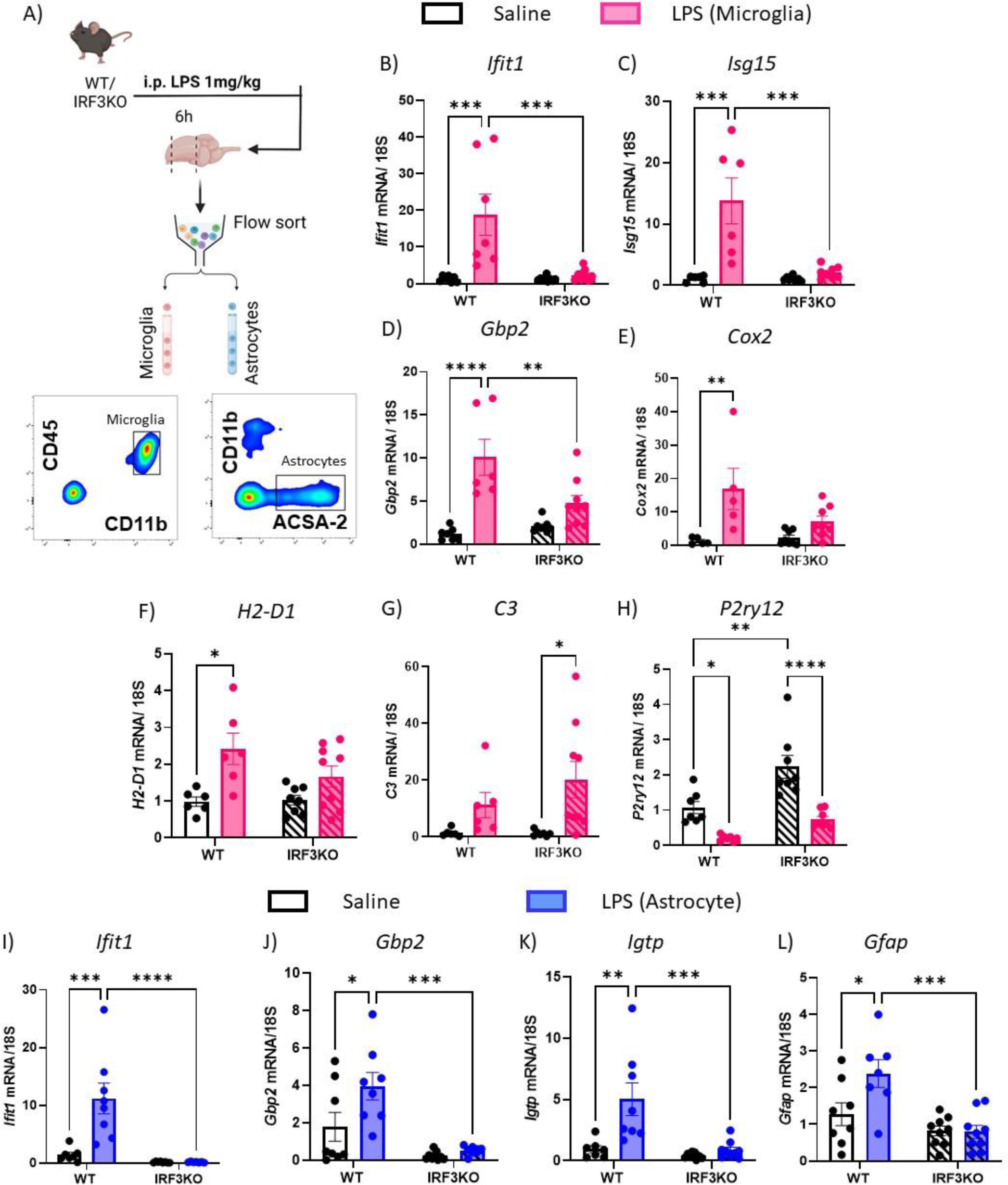
Transcripts from microglia & astrocytes of IRF3KO mice show a dampened proinflammatory response to LPS. A) Schematic of the protocol for isolation of microglia and astrocytes after acute LPS challenge. B-H) Quantification of qRT-PCR of transcripts from microglia show increased transcript levels of *Ifit1*, *Isg15, Gbp2, H2-D1, and Cox2* in WT-LPS group. While IRF3KO mice do not show significant induction of ISGs and certain proinflammatory transcripts in microglia (B-F), they appear more sensitive to LPS-mediated induction of *C3* transcripts (G). Also, compared to WT, IRF3KO microglia show a similar reduction in levels of *P2ry12* (H). N=6,9 per group. I-L) Quantification of qRT-PCR of transcripts *(Ifit1, Gbp2, Igtp, Gfap)* from astrocytes shows the attenuated response to LPS-induced transcripts compared to the WT controls. N=7,9 per group. Two-way ANOVA with Tukey’s multiple comparisons. *p<0.05, **p<0.01, ***p<0.001, ****p<0.0001

IRF3 is expressed by all the major cell types in the brain including astrocytes ^26^. Thus, we also tested the proinflammatory state of flow-sorted CD11b^-^ACSA-2^+^ astrocytes in IRF3KO mice (Fig 2A). We observed that LPS-induced upregulation of interferon signaling (*Ifit, Gbp2, and Igtp* mRNA) (Fig2 I-K) and *Gfap* (Fig 2L) were significantly lower in the IRF3KO-LPS mice compared to the WT-LPS group, suggesting that IRF3 is important for LPS-mediated astrocyte activation (Fig 2I-L).

### IRF3KO mice show reduced myeloid cell infiltration and inflammasome activation in the brain after repeated LPS challenges

IFN-I signaling is implicated in myeloid cell infiltration ^27^. However, the contribution of IRF3 specifically in the context of myeloid cell infiltration in the CNS is unexplored.

No monocyte infiltration was detected in response to 6h of single LPS injection in vivo in our model (Supplementary Fig 1A). Also, in chronic neuroinflammatory conditions TLR activation occurs constitutively or repeatedly. Thus, we next tested IRF3 activation and its downstream effects in a repeated LPS challenge paradigm. Here, mice were treated with a 1mg/kg dose of LPS daily for 4 days and euthanized 6h after the last LPS dose (Fig 3A).

**Figure 3:**
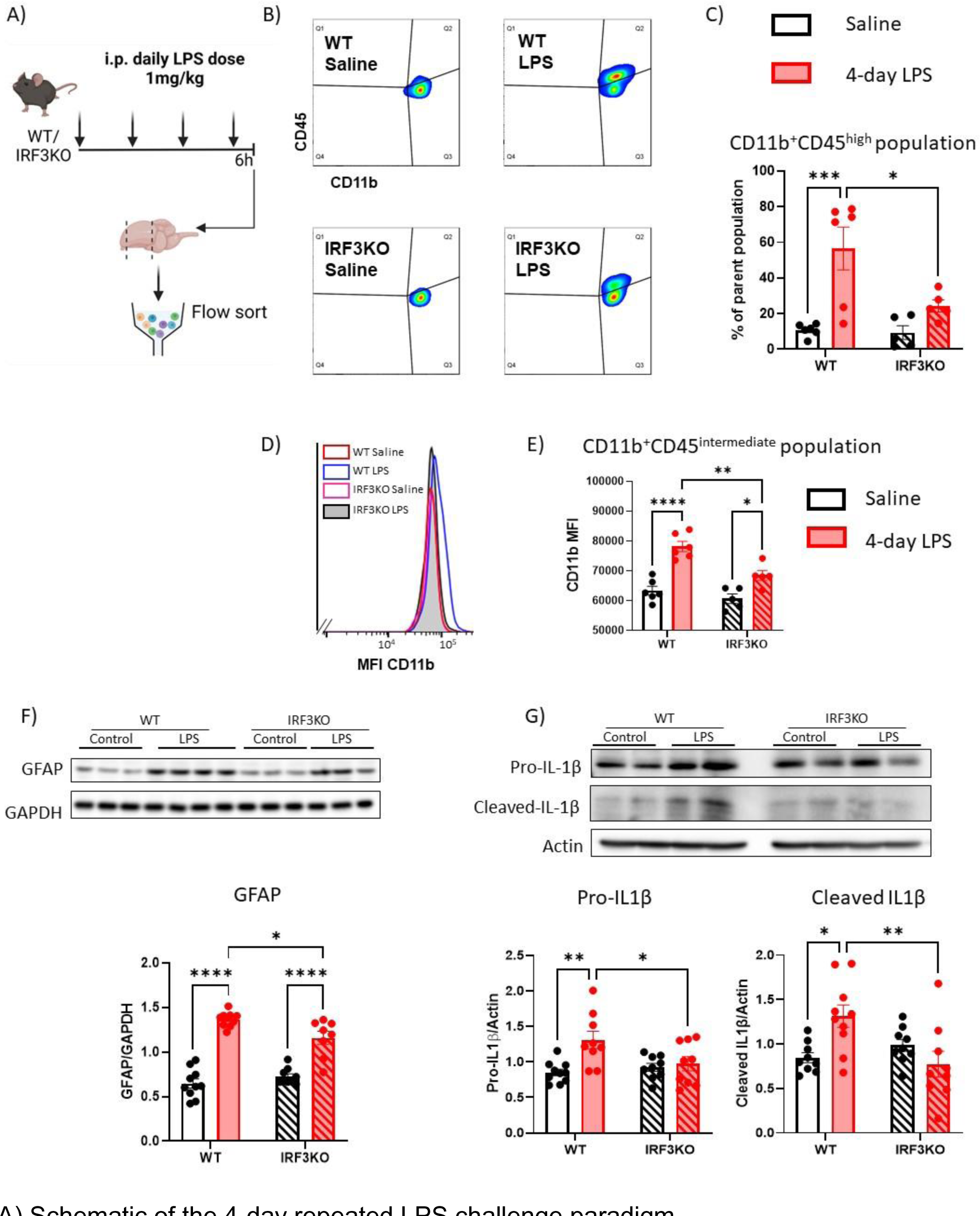
IRF3KO mice show reduced proinflammatory changes in the brain after repeated LPS challenges. A) Schematic of the 4-day repeated LPS challenge paradigm. B) Representative images of FACS analysis showing presence of significantly more infiltrating myeloid cells in Quadrant 2 (Q2)(CD11b^+^,CD45^high^) in the WT-LPS group in addition to the microglia population (CD11b^+^,CD45 ^intermediate^) in Q3. C) Quantification of the % of infiltrating cells shows that LPS-induced infiltration of myeloid cells was markedly reduced in the IRF3KO-LPS mice compared to the WT-LPS. N=5,6 each group. The parent population is defined as live cells based on DAPI staining. D-E) Quantification of the levels of mean fluorescence intensity of CD11b gated on the microglia in Q3 shows a more significant increase in WT-LPS microglia compared to IRF3KO-LPS microglia. N=5,6 each group. F) Representative images of western blots showing increased astrocyte proliferation in LPS-treated WT and IRF3KO samples in the cortical lysates. Quantification shows that the extent of astrocyte proliferation is significantly lower in IRF3KO mice compared to the WT. N=9,10 each group. G) Representative images and quantification of western blots showing a significant increase in the hippocampi of pro-IL1β (full length) and cleaved-IL1β indicate activation of inflammasome in LPS-treated WT samples. Quantification shows that IRF3KO mice are protected from this increase. N=9,10 each group. Two-way ANOVA with Tukey’s multiple comparisons. *p<0.05, **p<0.01, ***p<0.001, ****p<0.0001

In the WT-LPS group, we observed a distinct population of CD11b^+^,CD45^high^ cells, in addition to the resident microglia population defined as CD11b^+^,CD45^intermediate^, suggesting infiltration of peripheral myeloid cells upon repeated LPS challenges (Fig 3B,C). In contrast to WT, IRF3KO mice showed significantly reduced percentage of infiltrating myeloid cells (Fig 3B, C). We also determined that this myeloid cell infiltration took place in the absence of damage to the blood-brain barriers in our model of 4-day LPS challenge as indicated by no changes in the expression of blood brain barrier markers, Claudin-1 and Occludin (Supplementary Fig 2A-C).

Moreover, the microglia population of IRF3KO-LPS group showed significantly lower CD11b expression (gated on the CD11b^+^,CD45^intermediate^ microglia population) compared to the WT-LPS group (Fig 3D,E), further suggesting overall less proinflammatory effect of IRF3 deletion on microglia.

Because in the acute model we also observed proinflammatory transcripts in astrocytes, we tested whether astrocyte reactivity was also affected after 4 day repeated LPS challenge in IRF3 deficient mice. Assessment of GFAP levels in the cortex by western blots revealed a modest, yet significant, attenuation of GFAP levels in the IRF3KO-LPS mice compared to the WT-LPS mice (Fig 3F).

Because, we found attenuated IL1ß induction in the cortex after acute LPS challenge in IRF3KO mice (Fig 1C) we were curious to see if IRF3 contributed to inflammasome priming and activation. The effect of IRF3 perturbations on IL1β induction and inflammasome activation have not been tested in the CNS. Surprisingly, we could not detect the hallmark features of inflammasome activation in the cortical samples of the acute LPS or 4-day LPS challenged WT mice (Supplementary Fig 1B,C). LPS-mediated inflammasome activation has also been reported before in the hippocampus ^28^. Therefore we evaluated hippocampal lysates of the 4-day LPS challenged mice for inflammasome activation. Indeed we found, increased levels of pro- and cleaved-IL1β indicating inflammasome priming and activation in the WT-LPS group compared to the WT-saline mice. This increase in pro- and cleaved-IL1β levels was significantly attenuated in the IRF3KO-LPS group compared to the WT-LPS group (Fig 3G). This data revealed a novel role of IRF3 in inflammasome activation in the CNS as well as in regional sensitivity to LPS-mediated inflammasome activation.

Together, this data complements the observations in our acute LPS model and suggests that IRF3 deletion provides protection against various proinflammatory features of repeated LPS challenges such as myeloid cell infiltration, astrocyte proliferation, and inflammasome activation.

### IRF3 deletion delays TLR4 signaling and dampens cytokine secretion in primary microglia cultures

To investigate the specific role of IRF3 in microglia, we assessed how IRF3 modulates LPS-induced TLR4 signaling cascade. TLR4 activation leads to MyD88 dependent and independent signaling cascade that can feedback onto each other ^21, 29, 30^. To this end, we generated primary microglia cultures from WT and IRF3KO mice and challenged them in vitro with 20ng/ml LPS for various time points. We assessed phosphorylation of the key signaling cascades downstream of TLR4 activation: NF-κB (p65), p38, and ERK1/2 (Fig 4A).

**Figure 4:**
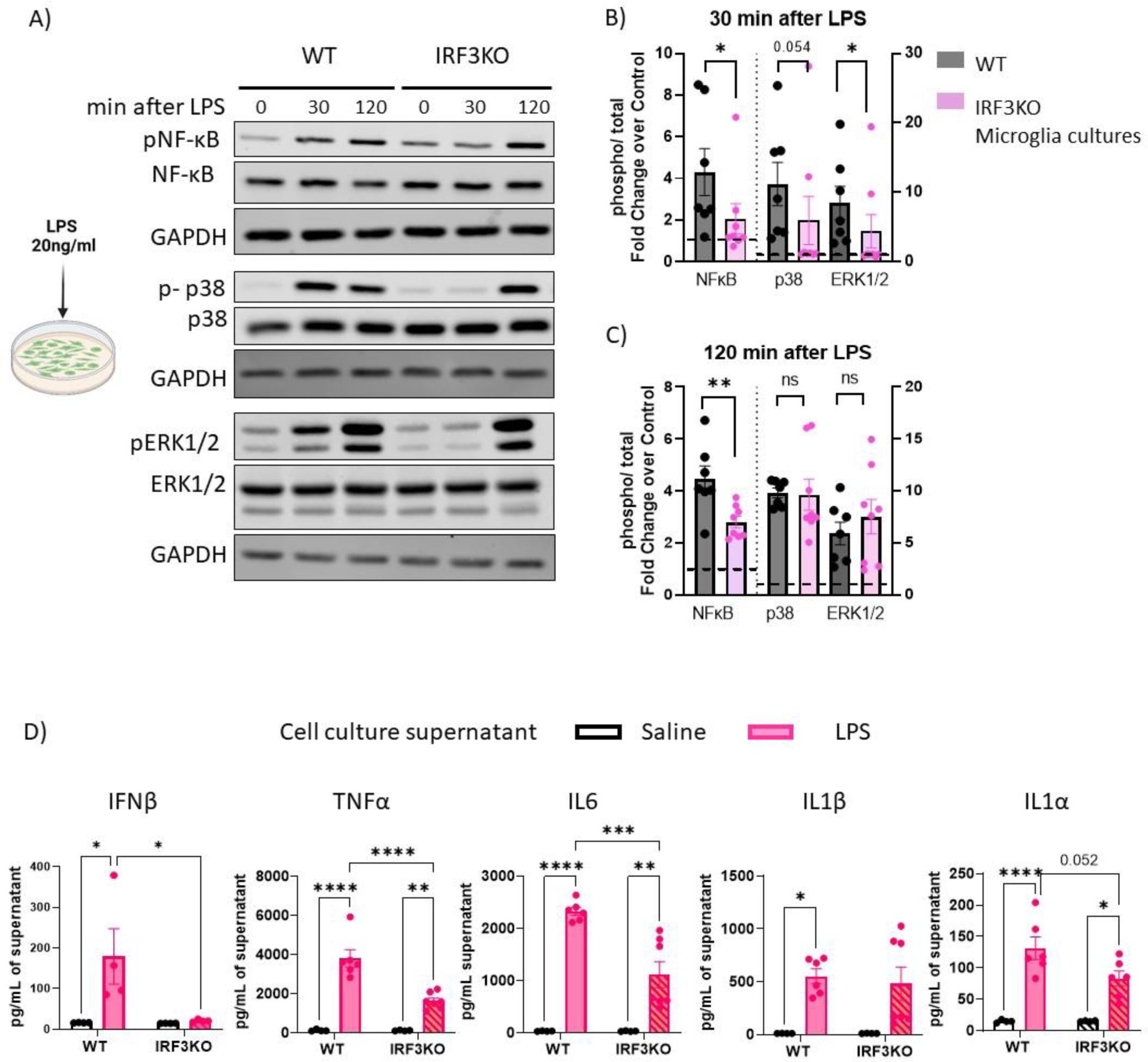
IRF3 deficient primary microglia cultures show delayed downstream signaling and attenuated cytokine production on LPS challenge. A) Representative images of western blots from primary microglia cultures treated with LPS. B) Quantification shows that 30 min after LPS stimulation, there is an increase in phosphorylation of NF-κB, p38, and ERK1/2 as indicated by the fold change over control of >1 in WT and IRF3KO cultures. However, IRF3KO microglia cultures show significant reduction in the levels of phosphorylation compared to that of WT. N= 7,8 for each group. Mann-Whitney test, *p<0.05, **p<0.01, ***p<0.001, ****p<0.0001. C) Quantification shows that 120 min after LPS stimulation, there is an increase in phosphorylation of NF-κB, p38, and ERK1/2 as indicated by the fold change over control >1 for both genotypes. However, IRF3KO microglia cultures only show a significant reduction in the levels of phosphorylation of NF-κB, while those of p38 and ERK1/2 are indistinguishable from that of the WT. N= 7,8 for each group. Mann-Whitney test or unpaired t-test as appropriate. *p<0.05, **p<0.01, ***p<0.001, ****p<0.0001. D) Quantification of Elisa from cell culture supernatants shows that proinflammatory cytokines (IFNβ, TNFα, IL6, ILβ, IL1α) are significantly upregulated in WT microglia on LPS challenge. IRF3KO cultures show either no release or significantly reduced release of cytokines on LPS challenge. N=4-7 for each group. Two-way ANOVA with Tukey’s multiple comparisons. *p<0.05, **p<0.01, ***p<0.001, ****p<0.0001

As expected, 30 min after in-vitro LPS addition, there was significant phosphorylation of the secondary signaling molecules-NF-κB (p65), p38, and ERK1/2 in the WT and IRF3KO microglia, as shown by the mean fold change >1 (over vehicle-treated samples) for phospho/total protein (Fig 4A, B). However, microglia isolated from IRF3KO mice showed strikingly lower phosphorylation levels of all the three signaling molecules at 30 minutes. Notably, the phosphorylation of NF-κB continued to be significantly lower in the IRF3KO cultures for up to 120 min after LPS stimulation whereas p-p38 and pERK1/2 were comparable to WT microglia (Fig 4A, C).

In addition, we observed significantly attenuated induction of cytokines in the supernatant of LPS-treated cultures of IRF3KO microglia (IFNβ, TNFα, IL6, and IL1α) when compared to the WT microglia (Fig 4D).

Together, our in vitro data shows an important regulatory role of IRF3 in LPS-mediated TLR4 signaling and cytokine production in microglia.

### Expression of a constitutively active form of IRF3 is sufficient to induce neuroinflammation

Phosphorylation of two serine residues (S388/390) is critical for IRF3 activation and nuclear translocation ^19^. Previously a constitutively active form of IRF3 i.e. IRF3-2D (S388D/S390D) was shown to induce proinflammatory cascade in macrophages and adipocytes ^23^. Thus, to specifically determine the effects of IRF3 activation in microglia, we expressed IRF3-2D in microglia using Cx3cr1Cre^ERT2^ and IRF3-2D-Lox mice.

We confirmed the expression of IRF3-2D constructs in EFYP+ cells from the brain at the transcript and protein levels (Supplementary Fig 3A). We observed the characteristic protein doublet for IRF3-2D in EYFP^+^ cells ^23^.

Similar to previous reports with Cx3cr1Cre^ERT2^ mice, we observed leaky expression of IRF3-2D in the absence of tamoxifen and a strong trend in further increase with tamoxifen administration (supplementary Fig 3A, B) ^31^. Therefore, we have also included additional Cre_only controls (Cre_Tam /Oil). Cre_Tam and Cre_Oil groups were very similar and thus data is pooled as a single group referred to as Cre_only.

Tmem119^+^ microglia from tamoxifen-administered IRF3-2D_Cre group (hereafter referred as IRF3-2D,Cre_Tam) (Fig 5A-D) showed significant morphological changes with reduced branching and intersections, compared to the IRF3-2D,Cre_Oil or Cre_only group suggestive of a reactive microglia morphology (Fig 5A,C,D). The morphological changes in the branching did not lead to the changes in cell volume (Fig 5B).

**Figure 5:**
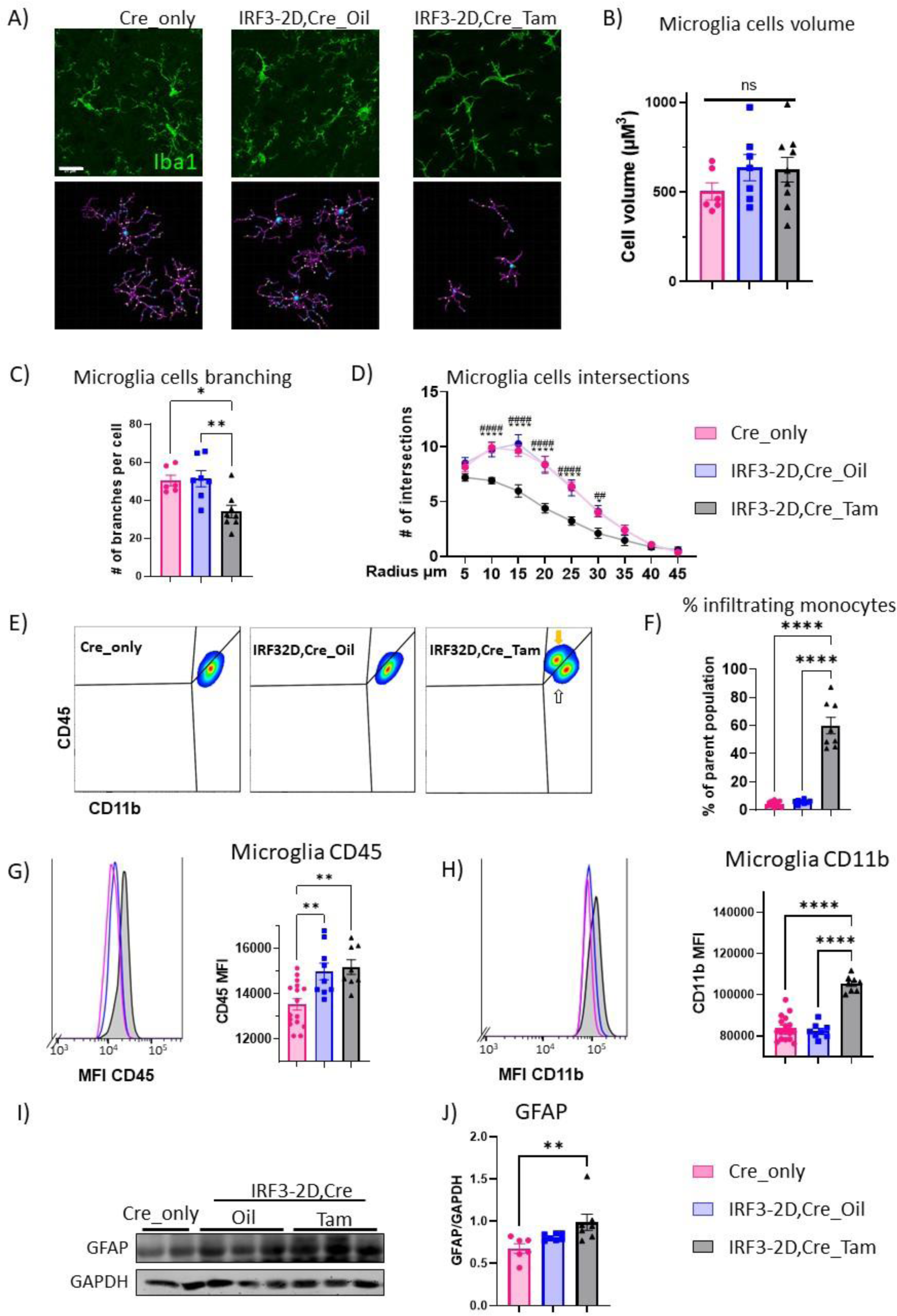
Expression of a constitutively active form of IRF3 is sufficient to induce neuroinflammation. A) Representative images of microglia (Tmem119^+^ cells were picked) co-stained with Iba1 and Scholl analysis performed using filament tracer software from Imaris. The scale bar is 21μM. B) Total volume of the cells did not change between any of the groups tested. C) Quantification of the microglia morphology shows that microglia (Cells positive for Tmem119) from IRF3-2D,Cre_Tam group show significantly reduced branching compared to IRF3-2D,Cre_Oil and Cre_only groups. D) Microglia from IRF3-2D,Cre_Tam group show reduced number of intersections at 10-30μm compared to that of IRF3-2D,Cre_Oil and Cre_only groups. * represents comparison with Cre_only, # represent comparison with IRF3-2D,Cre_Oil. For Scholl analysis N=6-8 for each group. >7 microglia were analyzed per animal. Data was analyzed using One-way ANOVA. *p<0.05, **p<0.01, ***p<0.001, ****p<0.000. E-F) Representative images and quantification of FACS analysis showing the presence of infiltrating myeloid cells (CD11b^+^,CD45^high^) in the IRF3-2D,Cre_Tam (Yellow arrow) group in addition to the microglia population (CD11b^+^,CD45^intermediate^) (White arrow). N= 8,9 each group. One-way Anova with Sidak’s multiple comparison test. ****p<0.0001 G-H) Quantification of CD45 and CD11b MFI gated on microglia (White arrow in E) shows significant upregulation in IRF3-2D,Cre_Tam, suggesting more reactive state compared to Cre_only controls. N= 8,9 each group. One-way Anova with Sidak’s multiple comparison test. ****p<0.0001 I) Quantification of the western blots shows proliferation of astrocytes, as measured by GFAP levels in cortical lysates, of IRF3-2D,Cre_Tam compared to Cre_only. N= 6,7 each group. Kruskal-Wallis test with Dunn’s multiple comparison test.

Moreover, flow cytometry of IRF3-2D,Cre_Tam brain samples revealed a distinct EYFP^+^ CD11b^+^CD45^high^ population of infiltrating monocytes in addition to Cd11b^+^CD45^intermediate^ microglia population (Fig 5E,F). This data corroborates the critical role of IRF3 in LPS-induced myeloid cell infiltration in the brain discussed earlier (Fig 3B,C). Moreover, the expression levels of CD45 and CD11b were elevated in the microglia population (gated on the Cd11b^+^CD45^intermediate^) of IRF3-2D,Cre_Tam mice compared to that of Cre_only mice (Fig 5G,H) further validating the proinflammatory microglia phenotype of IRF3-2D,Cre_Tam group.

Additionally, IRF3-2D,Cre_Tam mice also showed astrocyte reactivity in the cortex, suggesting that IRF3 activation is sufficient to mediate astrocyte reactivity (Fig 5I,J).

Taken together, these results demonstrate a proinflammatory role of IRF3 in microglia and astrocytes.

Despite this evidence of neuroinflammation, we found no significant behavioral changes in either of the anxiety tests (i.e. open field test and elevated plus maze) nor the Y-maze test, (Supplementary Fig. 4A-C) in the IRF3-2D,Cre_Tam mice compared to IRF32D,Cre_Oil or Cre_only group.

To gain deeper insights into the proinflammatory profile of IRF3-2D expressing cells, we performed bulk-RNA sequencing on flow sorted Cx3cr1^+^(EYFP^+^) population of myeloid cells from the brain (cortex, subcortical areas, and hippocampus).

To account for changes induced by tamoxifen administration, we compared the transcriptome of IRF3-2D,Cre_Tam EYFP^+^ population with that of Cre_Tam. We observed in total 908 genes that were differentially regulated in response to the presence of IRF3-2D. The expression of IRF3-2D in microglia resulted in a proinflammatory transcriptome enriched with the pathways related to IFN-β, IFN-𝞬, and viral responses (Fig 6A, B). In addition, we observed upregulation of pathways related to leukocyte migration, and cell adhesion further strengthening the effect of IRF3 on myeloid cell infiltration observed in this study (Fig 3B,C & 5E,F). We also found upregulation of pathways related to antigen presentation and co-stimulatory molecules [*H2 (-Ab1,-Eb1, -Aa, -Q6, -Q7, -K1, -D1, -Q5, -M3, -Dma,- K2, -T22, -Q4), Tap1, Cd74, Cd40, Cd72*), immunoproteasome (*Psmb9, Psme1, Psme 2*), cytoskeletal reorganization and ER-phagosome, providing further insights into the proinflammatory role of IRF3.

**Figure 6:**
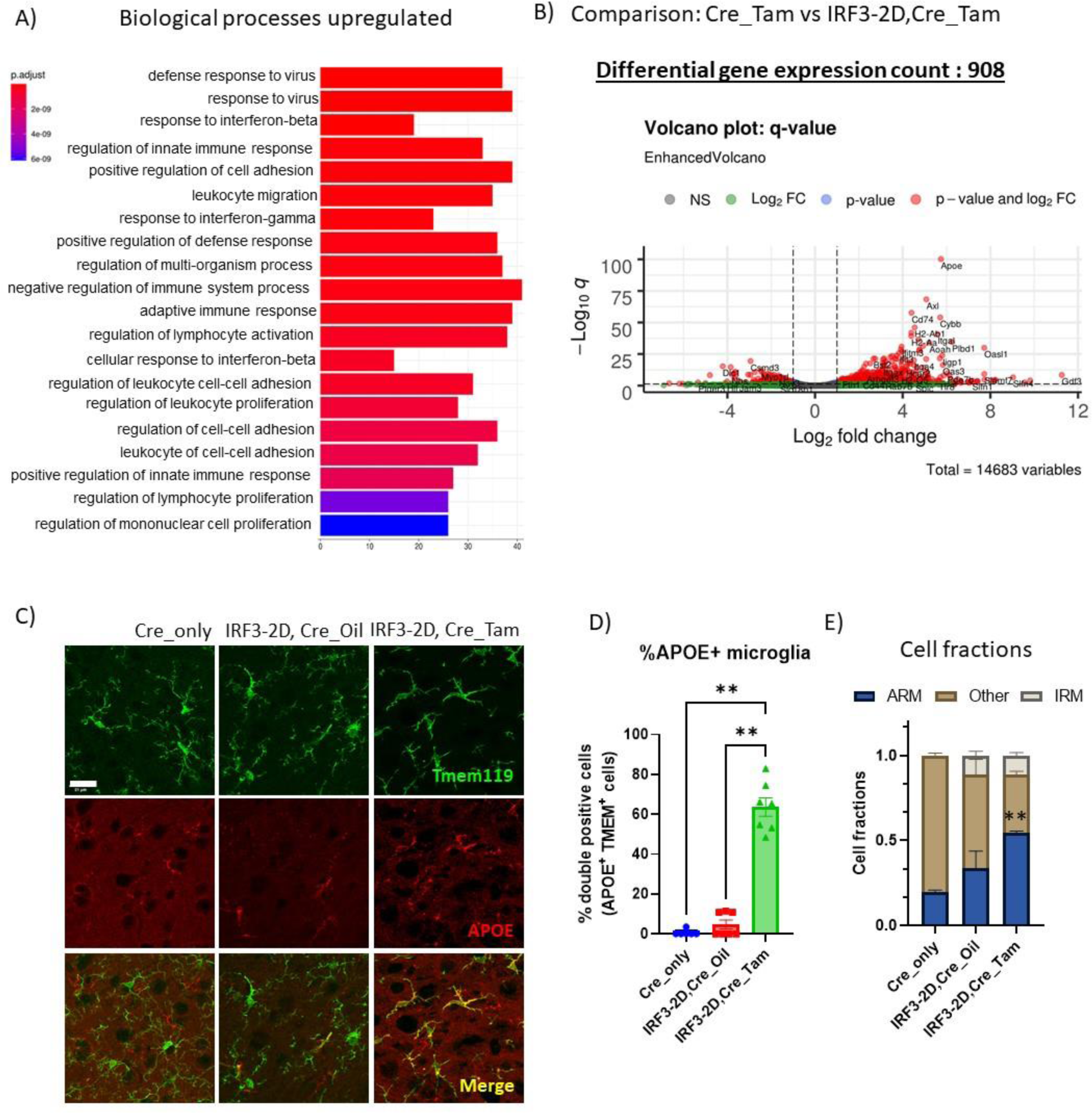
Overexpression of a constitutively active form of IRF3 leads to proinflammatory phenotype and induces expression of the AD risk genes. A) GO analysis of the differentially upregulated genes in FACS-sorted myeloid cells from IRF3-2D,Cre_Tam mice compared to Cre_only show proinflammatory phenotypes and upregulation of pathways related to interferon-β, γ signaling, cell adhesion, and leukocyte proliferation. B) Volcano plot showing differentially expressed genes in IRF3-2D,Cre_Tam mice compared to Cre_Tam. Note the upregulation of AD-associated genes. (n=2 for Cre_Tam, n=3 for IRF3-2D,Cre_Tam) C-D) Representative images of APOE staining in the cortex. Quantification confirms that APOE levels are significantly upregulated in the microglia from IRF3-2D,Cre_Tam group. The scale bar is 21μM. N= 6,8 per group. One-way Anova with Sidak’s multiple comparison test. ****p<0.0001. E) Deconvolution analysis on myeloid cells from IRF3-2D,Cre_Tam mice shows significantly more ARM-like cell fraction compared to Cre_only fraction. Both IRF3-2D,Cre_Oil and IRF3-2D,Cre_Tam cells contain IRM-populations in response to IRF3-2D-mediated signaling.

The top differentially regulated genes in this comparison were the subset of genes associated with AD. These included genes such as *Apoe, Axl, Cd74, Fth1, Itgax, and Ctsb* (Fig 6B). *As Apoe*, was the top candidate, we validated its expression at the protein level. We observed that APOE expression was significantly upregulated in Tmem119^+^ microglia in IRF3-2D,Cre_Tam group compared to IRF3-2D,Cre_Oil or Cre_only group (Fig 6C,D).

Microglia from the AD and neurodegenerative models show particular gene signatures which are termed as activated response microglia (ARM), or disease associated microglia (DAM) or microglia neurodegenerative phenotype (MGnD) with overlapping features ^14, 32–34^. In addition, interferon responsive microglia i.e. IRMs have also been reported in AD and aged mouse brains ^14, 16^. Therefore, we wondered what proportion of EYFP^+^ cells from IRF3-2D animals showed gene signatures associated with IRMs and AD. We performed data deconvolution with single cell RNA seq data to determine the cell fractions in IRM and ARM-like cells ^14^. The presence of IRMs was observed in IRF3-2D,Cre_Tam mice (Fig 6E) in line with the increased interferon signaling observed (Fig 6A). Interestingly, we observed significantly increased population of ARM in IRF3-2D,Cre_Tam group compared to the Cre_only group, and a strong trend in increase (p<0.09) compared to IRF3-2D_Oil group (Fig 6E). IRF3-2D,Cre_Oil group also showed presence of IRM and increasing trend in ARM population compared to the Cre_only controls, reflecting on the leaky expression of IRF3-2D and associated proinflammatory signaling in this model (Fig 6E, Supplementary Fig 5). Nonetheless, these results show that IRF3-mediated signaling is sufficient to induce IRM and ARM signatures in microglia. Thus, we conclude that IRF3 plays a critical role in microglia-mediated proinflammatory responses and regulates expression of genes associated with AD.

### Expression of ZBP1, a target of IRF3, is upregulated in microglia in various neuroinflammatory conditions

To further dissect the molecular mechanism and genes regulated by IRF3 signaling in microglia beyond LPS challenge or IRF3-2D model, we compared the transcriptome of IRF3-2D overexpressing EYFP^+^ cells to that of microglia from various neuroinflammatory conditions such as AD (5XFAD), Tauopathy model, LPS challenge, and glioma ^35^. In each data set, we used differentially upregulated genes showing a Log fold change of >0.6 and adjusted p-value of <0.05. From these comparisons, we identified 10 genes, comprising direct and indirect targets of IRF3, that are of relevance across different neuroinflammatory conditions (Fig 7A). IRF3-mediated changes in the transcriptome primarily result from the direct transcriptional activity of IRF3 or IRF3-mediated secondary signaling cascades. The direct transcriptional targets of IRF3 have been previously identified by ‘Cleavage Under Targets and Release Using Nuclease’ (CUT and RUN) technique from hepatocytes expressing IRF3-2D ^22^. Of these 10 common genes, 3 genes were identified as direct transcriptional targets of IRF3-*Oasl2, Zbp1 and Tlr2* by CUT and RUN ^22^.

**Figure 7:**
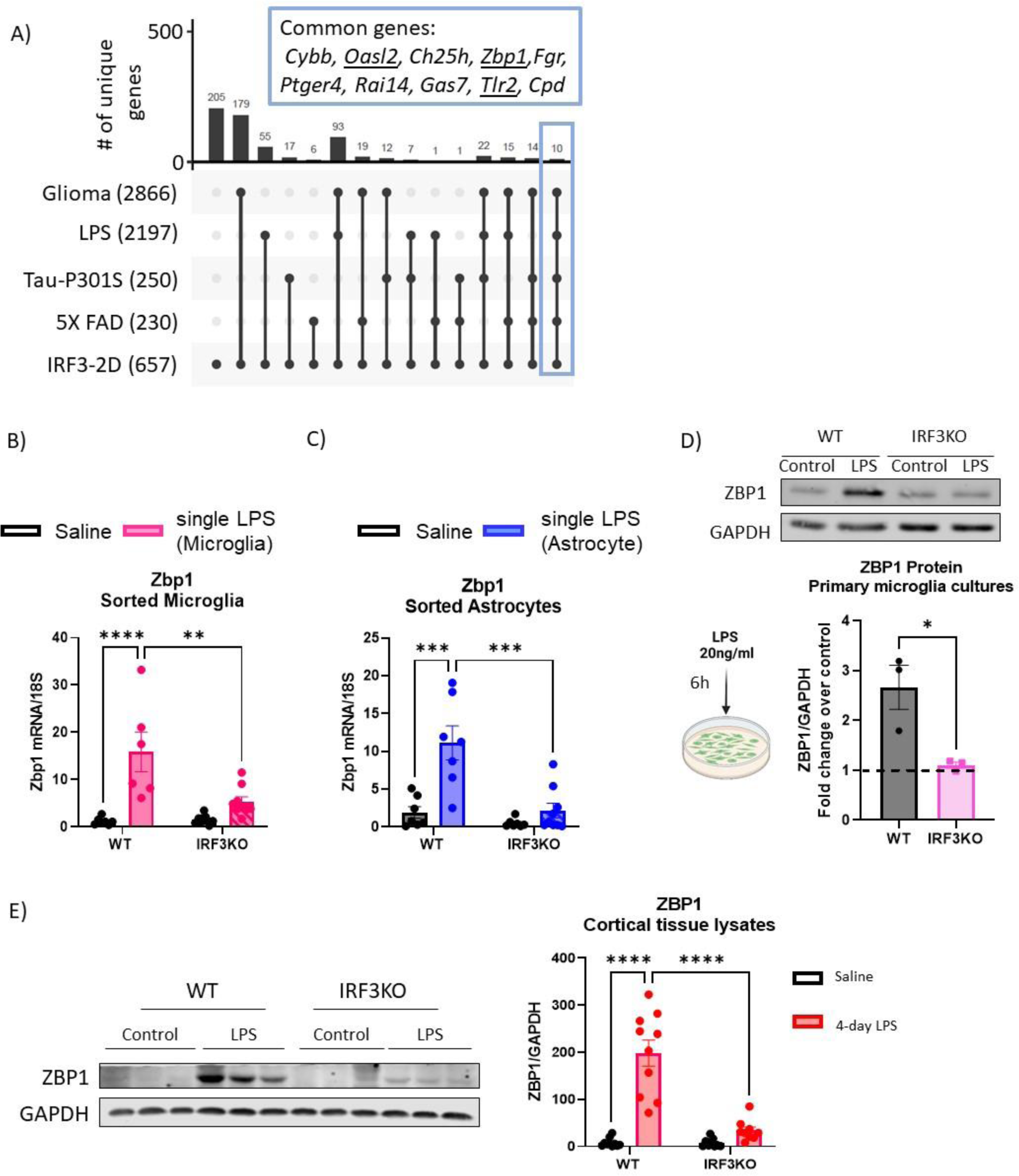
Zbp1 is a proinflammatory transcript common across various neuroinflammatory conditions and its expression is regulated by IRF3. A) An upset plot of differentially expressed genes in IRF3-2D expressing myeloid cells and microglia from various neuroinflammatory conditions. The number of differentially upregulated genes from each disease are represented in the bracket. Note the set of common genes across all five neuro-inflammatory conditions encased in blue. Underlined genes were identified as direct transcriptional targets of IRF3. B-C) Quantification of qRT-PCR of microglia and astrocytes sorted from acute LPS model (6h LPS challenge in vivo) shows upregulation of Zbp1 mRNA in WT, which is absent in IRF3KO condition. N=6-9 in each group. Two-way ANOVA with Tukey’s multiple comparisons. **p<0.01, ****p<0.0001 D) Representative image and quantification of western blot from microglia cultures treated with LPS for 6h show 2.6 fold induction in ZBP1 in WT microglia but not IRF3KO cultures. N=3 biological replicates. Unpaired t-test. *p<0.05 E) Western blot image and quantification of cortical tissue from WT and IRF3KO mice treated with LPS or saline for 4 days, show ZBP1 induction only in the WT-LPS group and absent in IRF3KO-LPS condition. N=9,10 each group. Two-way ANOVA with Tukey’s multiple comparisons. ****p<0.0001

In view of the novelty, we particularly focused on Zbp1. Zbp1 was initially recognized as interferon-inducible tumor-associated protein ^36^. ZBP-1 is shown to be critical for LPS-mediated production TNFα and IFNβ in macrophages ^37^. However, the role of ZBP1 in neurological disorders remains poorly studied. Thus, we aimed to validate Zbp1 as the target of IRF3 in our models of LPS challenge in the CNS.

We observed that after an in vivo acute LPS challenge *Zbp1* mRNA was significantly induced in microglia and astrocytes isolated from the WT-LPS group, while no change could be detected in cells isolated from the IRF3KO mice (Fig 7B,C). Similarly, there was a striking increase in the expression of Zbp1 (∼2.5 fold) in WT microglia cultures treated with LPS in-vitro when compared to IRF3KO primary microglia 6h after LPS stimulation (Fig 7D). This data corroborated results from the in vivo 4-day repeated LPS challenge model, where only the WT, and not the IRF3 deficient, brain tissue showed significantly elevated levels of Zbp1 protein after LPS treatment (Fig 7E).

Thus, together we identify Zbp1 as a novel proinflammatory target common across different neuroinflammatory conditions and show that Zbp1 expression is regulated by IRF3-induced signaling in microglia and astrocytes.

## Discussion

In this manuscript, we demonstrate that the expression of IRF3 in microglia is important in different neuroinflammatory contexts and IRF3 activation and IRF3-mediated signaling is sufficient to drive expression of AD-related genes. We also discovered that LPS-induced astrocyte activation is also dependent on IRF3. The function of IRF3 has been extensively studied in peripheral models of TLR3 and TLR4 activation i.e. viral and bacterial infection, respectively, including our work on IRF3 in sterile inflammatory conditions such as alcohol abuse and obesity ^23, 24, 38^. Role of IRF3 has been studied in viral encephalitis. Phosphorylation deficient mutation at S386 of IRF3 is associated with reduced IFN-I signaling in Herpes simplex encephalitis (HSE) patients ^39^. IRF3KO mice showed higher mortality rates and increase inflammation on HSE infection ^40^. Similarly, IRF3 deficient mice showed inability to resolve inflammation in the CNS by alphavirus infection ^41^. In this report we evaluated the cell type-specific contribution of IRF3 in microglia and its impact in a broader context of neuroinflammation.

Using LPS, a prototypical pathogen-associated molecular pattern (PAMP) and TLR4 ligand, we observed that IRF3 is involved in the production and release of the key inflammation-associated cytokines and chemokines, sickness behavior as well as Type 1 interferon-dependent genes in the brain. Other mediators, such as TNFα and IFNβ also contribute to sickness behavior, however, we could not detect a significant amount of these cytokines in the cortex of mice at 6h post LPS stimulation when other markers were assessed in our experiments ^42, 43^ .

Activation of TLR4, a widely studied pattern recognition receptor, has been observed in myriad of neuropathologies ranging from gram-negative bacterial infections (mimicked here by LPS), AD, Parkinson’s disease (PD), multiple sclerosis to amyotrophic lateral sclerosis ^44–47^. TLR4 also senses both pathogen-associated molecular patterns, such as LPS, and sterile inflammatory signals, for example HMGB1 ^48, 49^. Furthermore, LPS primes brain responsiveness to HMGB1 ^50^. Thus detailed understanding of TLR4 mediated downstream signaling in CNS is warranted. TLR4 triggers two downstream pathways through adaptor proteins: MyD88 and TRIF dependent leading to MyD88 independent signaling ^51^. While much attention is paid to MyD88-dependent or NF-κB-mediated signaling, here we highlight the role of IRF3 in TLR4/LPS-mediated inflammatory responses in neuroinflammation.

In addition to TLR4, IRF3 can also be activated intracellularly via the cGAS-STING pathway as well as via endoplasmic reticulum stress via STING ^38, 52, 53^. Microbial or endogenous DNA is recognized through cGAS-STING pathway and culminates in IRF3 activation ^52^. cGAS-STING activation is observed in various neuroinflammatory conditions such as AD, TBI, PD, aging etc. ^6, 7, 54^, indirectly implicating IRF3 in these conditions and emphasizing the need to study functions of IRF3 in the CNS.

In neuroinflammatory diseases there is continued presence of disease associated molecular patterns (DAMPs) and/or PAMPs that sustain inflammation. Thus, we also tested a four-day model of repeated LPS stimulations where we discovered a novel role of IRF3 in monocyte infiltration and inflammasome activation in the CNS. The IFN response in the CNS has been associated with myeloid cell infiltration under tumorigenic conditions and viral infections ^27, 55^, however, the specific role of IRF3 in myeloid cell infiltration in the brain has not been described previously. The reduced myeloid cell infiltration observed in the IRF3KO-LPS group in the 4-day model correlates with the reduced levels of MCP1 seen in the acute model in Fig 1C. Interestingly, at 4 day time point, we could not detect MCP1 anymore in the samples, suggesting that MCP1 release in the initial LPS challenge is sufficient to elicit myeloid cell infiltration in the brain.

In our study, reduced NLRP3 inflammasome activation modulated by IRF3 deletion in the CNS was another novel finding. This observation is significant in light of the critical role of NLRP3 in AD, and other neurological disorders ^56, 57^. This result is in line with the previous observations made by our lab and others showing reduced NLRP3 inflammasome activation in the absence of IRF3 in the peripheral models of inflammation^24, 58^.

IRF3 is expressed in the brain by microglia, astrocytes, neurons, endothelial cells and oligodendrocytes ^26^. As astrocytes have increasingly gained importance to partake in regulating immune responses in the brain, we assessed the responses from astrocytes. Indeed we found a significant reduction in the LPS-induced proinflammatory response of astrocytes in the absence of IRF3 expression in both the in vivo models of LPS that we tested, suggesting a major role for IRF3 in astrocyte responses to acute as well as repeated LPS challenges. Proinflammatory responses of microglia contribute to astrocyte activation, however, in our study we cannot distinguish between microglia dependent or astrocyte autonomous role of IRF3 in LPS-mediated astrocyte activation^17, 59^.

LPS-induced upregulation of proinflammatory transcripts in microglia of WT and IRF3KO mice, showed partial dependence on IRF3. Surprisingly, we observed complement factor *C3* transcripts were upregulated in IRF3KO-LPS microglia compared to WT-LPS groups. This finding is surprising since *C3* is a known target of IFN-I ^4, 60^ and is also significantly upregulated in IRF3-2D expressing myeloid cells (Fig 6B). A compensatory effect of LPS-mediated IRF3-independent pathway of complement activation may explain this effect ^61^. Moreover, microglia transcripts from IRF3KO mice showed elevated levels of the homeostatic marker *P2ry12* compared to the WT. However, solely based on this result it is difficult to draw conclusions on the homeostatic state of microglia in these mice. Analysis of the transcriptome of IRF3KO mice microglia may shed light on this aspect.

To determine the role of IRF3 in LPS-mediated signaling in microglia we used primary microglia cultures derived from WT and IRF3KO pups. As seen in the cortical lysates, LPS-induced cytokine release in the supernatant was reduced in IRF3KO microglial cultures compared to WT. Furthermore, we observed that IRF3 deletion directly modulated the signaling events downstream of TLR4 particularly in the first 30 min compared to 120 min, suggesting that effects of IRF3 deletion are compensated by the feedback loops between secondary messengers downstream of TLR4 signaling.

IRF3KO mice have been shown to have reduced peripheral inflammation in response to LPS ^25^. Our *in vitro* results showed reduced cytokine levels and proinflammatory responses of isolated IRF3 deficient microglia compared to wild type, indicating that the observed reduction in neuroinflammation in vivo in IRF3 KO mice was not just an outcome of reduced peripheral inflammation but also reduced intrinsic inflammatory response of IRF3KO microglia to LPS challenge.

While our data in IRF3KO mice and cells indicated the importance of this pathway in TLR4-mediated neuroinflammation, next, to understand the isolated effects of IRF3 activation in microglia we took advantage of the IRF3-2D-lox line, described previously^23^. Our data indicate that constitutive IRF3 activation in microglia results in key features of neuroinflammation including increased monocyte infiltration to the brain and increased GFAP expression suggesting astrocyte activation. We also found that a key feature of IRF3-2D expression was the upregulation of certain DAM genes, or ARMs, most notably *Apoe*. *Apoe* is a major risk factor for the late onset Alzheimer’s disease. In addition, *Apoe* expression in microglia has been shown to regulate microglia immunometabolism influencing their ability to respond to Aβ plaques, and tauopathy ^32, 62–64^. APOE-TREM2 pathway has been shown to be important for expression of DAM and ARM genes ^14, 32^. Our model of IRF3-2D, suggests that sustained IRF3 activation is sufficient to drive the expression of *Apoe*, which in turn can regulate the expression of certain genes associated with microglia phenotype in neurodegenerative diseases. *Apoe* is not one of the known transcriptional targets of IRF3; our study suggests that it may be upregulated through IRF3-mediated mechanisms. Further investigation is needed to determine the exact mechanism of IRF3-mediated upregulation of APOE.

Since the bulk RNAseq performed from IRF3-2D,Cre_Tam mice comprises EYFP^+^ cells in the brain i.e. microglia and infiltrating myeloid cells, we ascertained microglia specific effects by visualizing Tmem119^+^ cells for morphological analysis (Fig 5A), and APOE expression (Fig 6C) and using CD11b^+^CD45^intermedate^ gate for assessing levels of CD11b in Fig 5G,H). We also compared the transcriptome of EYFP^+^ cells devoid of infiltrating myeloid cells from IRF3-2D,Cre_Oil with Cre_Oil groups (Fig 5E,F and Supplementary Fig 5). This comparison showed a total of 321 differentially regulated genes (DEGs), fewer than the 908 DEGs described in IRF3-2D,Cre_Tam group in Fig 6. Here, we observed proinflammatory pathways and genes such as *Axl, Cybb, Cst7, H2-D1, Cd74* (Supplementary Fig 5) and increasing trend in ARM fraction (Fig 6E), collectively showing proinflammatory effect of IRF3-2D activation on microglia in the absence of infiltrating myeloid cells.

While these results clearly establish effects of IRF3-2D on microglia, we cannot rule out the effect of leaky expression of IRF3-2D in Cx3CR1^+^ myeloid cells in the periphery and further experiments would be needed to tease those apart. It is interesting to note that Tamoxifen administration further increased the ARM-like fraction in IRF3-2C,Cre_Tam group while the IRM-like fraction shows no additive effect, it is possible that the expression of ARM related genes is induced by targets of IRF3 not directly involved in IFN-I signaling (CUT and RUN analysis ^22^). Previously, IFN-I signaling in microglia has been associated with increased anxiety ^9, 65^, however despite induction of such a strong IFN-I signaling cascade in the IRF3-2D brains, we found no obvious behavioral changes in anxiety or memory performance of these mice. In our model we cannot rule out the development of any compensatory behavioral and transcriptional changes that may mask the subtle underlying behavioral abnormalities. Moreover, in this model we see IRF3-mediated signaling which may not recapitulate the full spectrum of inflammation and IFN-I signaling observed by others (Ben-Yehuda et al., 2020, Sahasrabuddhe and Ghosh, 2022).

Lastly, to evaluate the presence of signatures of IRF3 activation and IFN-I signaling in different proinflammatory disorders, we compared the genes upregulated with IRF3-2D expression to that of the genes upregulated in different neuroinflammatory disorders such as glioma, Alzheimer’s disease model of amyloid and tauopathy, and LPS challenge. Of these 10 common genes, we were particularly interested in Zbp1. Zbp1 is known for its function in cell death pathways, viral response and inflammasome activation ^36^. In addition, the role of Zbp1 in proinflammatory signaling, independent of cell death, is also emerging ^66^. However, there are limited studies investigating the role of Zbp1 in neuroinflammation and its role in AD is beginning to emerge ^67^.

Previous studies have shown Zbp1 to be a regulator of IRF3 ^37^. We recently showed that Zbp1 expression is modulated by IRF3 in mouse models of cholestatic-liver injury ^24^. Here we show for the first time that IRF3 can directly regulate Zbp1 levels in microglia and astrocytes.

Taken together we discovered new insights into the role of IRF3 in promoting neuroinflammation specifically, in microglia and highlight IRF3 and its downstream genes as important players in various neuroinflammatory conditions.

## STAR Methods

### Mice

The following mice were used-C57BL/6 from Jax mice (000664), IRF3KO (described previously, ^24^, Cx3cr1^CreERT2^(B6.129P2(Cg)-Cx3cr1tm2.1(cre/ERT2)Litt/WganJ-021160), IRF3-2D (C57BL/6-Gt(ROSA)26Sortm4(CAG-Irf3*S388D*S390D)Evdr/J-036261). All strains were in C57BL/6J background. The mice were maintained on ad-libitum food and water. All the breedings, experiments and euthanasia were conducted as per the institutional IACUC protocol 030-2022. Both the sexes between the ages of 3-6 months were used.

### Tamoxifen preparation

Tamoxifen stocks of 20mg/ml were prepared by dissolving Tamoxifen in Corn oil at 37^0^C. Mice were given oral gavage 10mg/kg of Tamoxifen or equal volume oil for consecutive 5 days and used for experiment 5 weeks later.

### LPS preparation and administration

LPS was prepared by dissolving LPS in saline at 1mg/mL and intraperitoneally injected in mice at 1mg/kg dose as indicated. For in vitro experiments LPS was dissolved in water at 100ug/mL concentration and diluted in media just before addition.

### Microglia and astrocyte flow cytometry

Microglia and astrocytes were flow sorted as described previously ^68^. Briefly, mice were transcardially perfused and brains were dissected out. One half of the brains were fixed in 4%PFA overnight. From the other half the prefrontal cortex, hippocampus and cerebellum were dissected out and frozen on dry ice. The rest of the brain was homogenized in ice-cold HBSS (Ca^++^, Mg^++^ free). Cells were pelleted at 350g for 7 min followed by a 37% percoll plus spin without brakes. The top layer of myelin was aspirated and the microglia pellet was washed in HBSS before staining. The cell pellet was incubated in FC block (1:50) at 4^0^C for 5 min followed by incubation in antibodies against CD11b, CD45, and ACSA-2 in FACS buffer (2% Fetal Bovine Serum in PBS Ca^++^, Mg^++^ free) at 4^0^C for 20 min. DAPI (1mg/ml, 1:1000) was added in the last 5 min of antibody incubation. The cells were washed in FACS buffer and sorted using Cytoflex-SRT or analyzed on Cytek Aurora. Microglia were sorted as Cd11b^+^, CD45^intermediate^ population and astrocytes were sorted as CD11b^-^, ACSA-2. For mice in Cx3cr1^CreERT2^ background cells were sorted using EYFP fluorescence. Sorted cells were pelleted and stored at -80^0^C until downstream processing. 15-20K microglia were used for western blotting. Flow data was analyzed using FlowJo.

### Primary microglia cultures

Primary microglia were cultured as described previously with slight modification ^69^. Brains from the WT and IRF3KO pups (0-4 days old) were dissected, meninges removed, and homogenized with mortar and pestle. Cells were pelted by centrifugation at 350g for 7min at 4^0^C and directly plated onto Poly-D-Lysine (PDL) coated (10ug/mL) 90mm dishes. Cells were cultured in DMEMF-12 containing 10% FBS and 1% Penicillin/Streptomycin. Cultures were grown at a standard 5% CO2, 37^0^C incubator. Next day cultures were washed 3 times with phosphate-buffered saline and incubated for an additional 3-4 days in the culture medium described above before the addition of the growth factors (mCSF and TGFβ). 2-3 days later microglia were shaken off the astrocyte monolayer and harvested every 3rd day for 3 cycles. Harvested microglia were plated on PDL coated 12 well dish at 4 x 10^5 cells per/mL in plain DMEM-F/12 without FBS a day before the experiment. On the day of the experiment, cells were treated with LPS (20ng/mL) for an indicated amount of time, and supernatant and cells were harvested for further analysis. The supernatant was spun at 10K for 10 min at 4^0^C and stored at -80^0^C until further use. Cells in each well were washed in ice-cold PBS before harvesting.

### Western Blotting

RIPA was used as a lysis buffer with a Protease and Phosphatase inhibitor cocktail. Brain tissue was lysed in the tissue homogenizer, followed by a spin at 10K for 10 min at 4^0^C. A predetermined number of cells as indicated above was loaded for western blots from primary microglia cultures or flow-sorted microglia. Total of 50ug of protein was loaded onto SDS gels from tissues. Proteins were transferred onto nitrocellulose membranes and blocked in 5% BSA in 0.1% TBST at room temperature (RT) for 1h. Blocked membranes were incubated with primary antibodies in 5% BSA overnight and washed 3 times in 0.1% TBST. A secondary antibody was added in blocking solution for 1h at RT followed by 3 washes in 0.1% TBST before developing the blot.

For western blots from primary microglia in Figure 4B & C, the results are represented as a comparison between WT and IRF3KO cultures. For signaling cascades the phosphorylation levels are represented as = (LPS-treated [Phospho protein/ Total protein]) / (Saline-treated [Phospho protein/ Total protein]) for WT and IRF3KO cultures separately.

### Immunohistochemistry and image analysis

Brains were fixed in 4% PFA overnight, followed by cryopreservation in 30% sucrose solution until the brains sank. Brains were sectioned using Leica cryostat into 25μm thin sections, collected in 0.05% Sodium Azide solution in PBS, and stored at 4^0^C until stained. Desired brains were mounted onto glass slides, washed in PBS and blocked using 1% Triton and 10% Horse Serum in PBS for 1h at RT. Primary antibodies were dissolved in the blocking and incubated overnight at 4^0^C. Primary antibodies were washed at RT in 1% Triton in PBS and incubated in secondary antibodies in 1% Triton and 1% Horse serum for 1h at RT. After secondary antibody incubation, DAPI (1mg/ml at 1:1000) solution was added for 5 min at RT followed by 2 washes with 1% Triton in PBS. Sections were imaged at 63x magnification on a Zeiss LSM-700 confocal microscope. Iba^+^ staining was used for morphometry analysis. In Fig 5A, the sections were co-stained with Tmem119, a microglia specific marker, to verify the microglial identity of the cells. The filament tracer module and Sholl analysis extension in Imaris (Bitplane; Zurich, Switzerland) were used to assess microglia morphometry.

### Bulk-RNA seq

1000 sorted microglia were suspended in 1% beta-mercaptoethanol in TCL buffer and sequenced using smart-seq2 platform at Broad Institute. Briefly, the raw sequencing reads were quality-checked and data were pre-processed with Cutadapt (v2.5) for adapter removal. Gene expression quantification was performed by aligning against the GRCm38 genome using STAR (v2.7.3a). Reads were quantified against Ensembl v98 annotated transcript loci with feature Counts (Subread 1.6.2). Differential gene expression analysis was performed using DESeq2 (v1.24.0) while ClusterProfiler (v3.12.0) was utilized for downstream functional investigations. Plots were generated in R using ggplot2 (v3.3.3), EnhancedVolcano (v1.8.0), ComplexHeatmap (v2.6.2.

For deconvolution analyses we reanalyzed previously published single-cell expression data as described in the original manuscript (GSE127884). The data containing labels for ARM and IRM cell types was uploaded to the CIBERSORTx platform in order to generate a signature matrix for these cell populations. This matrix was used in combination with our bulk RNA data in order to estimate the relative amounts of each of the cell types.

### RNA isolation and qRT PCR

RNA was isolated from microglia and astrocyte pellets using Qiagen RNeasy plus micro kit. cDNA was converted using the Superscript II kit. Gene expression analysis was conducted by qRT PCR using SYBR green from BioRad. Gene expression for every sample was normalized to 18s rRNA as housekeeping gene.

### Elisa

Cytokine levels were detected using Elisa kits. Plates were coated as per manufacturer’s instructions. Cell culture supernatant or tissue lysates (prepared as described above) were incubated overnight at 4^0^C. Kit-specific protocol was followed for washing and developing of the Elisa plate. Absorbance was measured on a microplate reader and the amount of the cytokine was estimated based on the standard curve.

### Animal behavior

Mice were brought into the behavior room 30 min before the experiment. Animal behavior was recorded for 5 min for all tests with an overhead camera and analyzed using Ethovision^XT^. 20 lux light intensity was maintained in the room. Animal behavior was conducted between 9am-2pm. The position of the animal was monitored using the center of mass body point. The behavior tests were performed at least 24h apart.

For an open field test, 40cm x 40cm x 40cm arena was used. Mice were released in the center of the arena, facing away from the experimenter. As indicated in the figures, the total distance traveled, velocity and time spent in the center were calculated to determine anxiety-like behavior or sickness behavior. The central zone was marked 5 cm away from the walls of the arena.

For the elevated plus maze, mice were released into the central zone facing the open arms away from the experimenter. Total time spent in the open arms was used as the measure of anxiety. The length of each arm was 20 cm.

Y maze test was used to assess memory performance in mice. The Y maze test was performed last, where mice were allowed to explore the arena, and memory was assessed based on the pattern of spontaneous arm alternation. The length of the arms was 15 cm each.

### Statistics

Data was plotted as mean ± SEM using GraphPad Prism 9. The appropriate statistical test is indicated in the figure legend for each comparison.

## Supporting information

Resource Table

## Funding

This study was supported by NIH grant-R01AG072899 and NIAAA grant-5R01AA017729.

### Acknowledgments

We would like to thank NIH and NIAAA grants for funding and support. Figures-schematics were created using BioRender.com. We would like to thank Dr. Evan Rosen for generous gift of IRF3-2D (C57BL/6-Gt(ROSA)26Sortm4(CAG-Irf3*S388D*S390D)Evdr/J-036261) mice.

