## Supplementary material for "IRF3 regulates neuroinflammatory responses and the expression of genes associated with Alzheimer’s disease": Resource Table

Key resource table:

| Reagent and resources | Source | Identifier |
| --- | --- | --- |
| **Antibodies:** |  |  |
| Anti-APOE | Cell Signaling Technologies | 49285 |
| APC anti-mouse/human CD11b [M1/70] | Biolegend | 107636 |
| Anti-CD45 Rat Monoclonal Antibody (Brilliant Violet® 785) [clone: 30-F11] | Biolegend | 123109 |
| Anti-Iba1 Rabbit Polyclonal Antibody | Wako Chemicals | 019-19741 |
| Anti- ACSA-2 Antibody, anti-mouse, PE | Miltenyi Biotec | 130-123-284 |
| Anti- Irf 3 (D83B9) Rabbit mAb | Cell Signaling Technologies | 4302 |
| Anti-GFAP antibody | Abcam | ab4674 |
| Anti- Iba1 polyclonal guinea pig antibody | Synaptic Systems | 234 308 |
| Anti-IL1b antibody | GeneTex | GTX74034 |
| Anti-Claudin-1 antibody | Cell Signaling Technologies | 4933 |
| Anti-Occludin antibody | Abcam | ab167161 |
| Anti pNF-kB antibody | Cell Signaling Technologies | 3033 |
| Anti NF-kB antibody | Cell Signaling Technologies | 8242 |
| Anti GAPDH antibody | Proteintech | 60004-1 |
| Anti p-P38 antibody | Cell Signaling Technologies | 4511 |
| Anti P38 antibody | Cell Signaling Technologies | 9212 |
| Anti p-ERK1/2 antibody | Cell Signaling Technologies | 4370 |
| Anti ERK1/2 antibody | Cell Signaling Technologies | 9102 |
| Anti-ZBP1, mAb (Zippy-1) | AdipoGen | AG20B0010C100 |
| Goat anti Guinea Pig Secondary Antibody, Alexa Fluor 647 | Thermo Scientific | A-21450 |
| Goat anti-Rabbit Secondary Antibody, Alexa Fluor 488 | Thermo Scientific | A11008 |
| Goat anti-Mouse Secondary Antibody, Alexa Fluor 546 | Thermo Scientific | A-11030 |
| Rat anti-mouse CD16/CD32 Fc Block | BD Pharmingen | 553142 |
| Poly-D-lysine hydrobromide,mol wt 70,000-150,000 | Sigma-Aldrich | P0899-10MG |
| **Softwares:** |  |  |
| EthoVision | Noldus | Ethovision XT |
| (Base license plus the Multiple Body Points Module) |  |  |
| FlowJo | FlowJo |  |
| ImageJ | National institute of Health |  |
| Imaris | Bitplane |  |
| **Reagents:** |  |  |
| Lipopolysacchrides | Sigma-Aldrich | L4391 |
| Tamoxifen | Sigma-Aldrich | T5648 |
| Percoll PLUS | GE Healthcare | 17–5445–02 |
| RNeasy Plus Micro Kit | Qiagen | 74034 |
| IL1-α Elisa Kit | R & D systems | DY400 |
| IL-1β Elisa Kit | R & D systems | MLB00C |
| MCP-1 ELISA MAX™ Deluxe ELISA Kit, | BioLegend | 432704 |
| Mouse IL-6 DuoSet ELISA | R & D systems | DY406 |
| Mouse CXCL1/KC DuoSet ELISA | R & D systems | DY453 |
| Mouse TNF-α DuoSet ELISA, | R & D systems | DY410 |
| IFNβ Elisa kit | R & D systems | MIFNB0 |
| DMEM/F-12, no phenol red | Thermo Scientific | 21041025 |
| Recombinant human TGF-b | PeproTech | 100-35 |
| mouse colony-stimulating factor-1 | Peprotech | 315–02 |
| Protease/phosphatase Inhibitor Cocktail (100X) | Cell Signaling Technology | 5872S |
| Applied Biosystems Power SYBR Green PCR Master Mix | Thermo Fisher Scientific | 43-687-02 |
| Iscript™ Reverse Transcription Supermix, | Bio-Rad | 1708841 |
| Normal horse serum | Vector Labs | S-2000-20 |
| cOmplete, Mini, EDTA-free Protease Inhibitor Cocktail | Roche | 4693159001 |
| **Primer sequences:** | Forward 5'-3' | Reverse 5'-3' |
| Ifit1/ isg-56 | GCCTATCGCCAAGATTTAGATGA | TTCTGGATTTAACCGGACAGC |
| isg-15 | GGTGTCCGTGACTAACTCCAT | TGGAAAGGGTAAGACCGTCCT |
| gbp2 | AGATGCCCACAGAAACCCTCCA | AAGGCATCTCGCTTGGCTACCA |
| cox-2 | GCTGTACAAGCAGTGGCAAA | CCCCAAAGATAGCATCTGGA |
| h2-d1 | TGAGGAACCTGCTCGGCTACTA | GGTCTTCGTTCAGGGCGATGTA |
| c3 | CGCAACGAACAGGTGGAGATCA | CTGGAAGTAGCGATTCTTGGCG |
| p2ry12 | CACCTCAGCCAATACCACCT | CAGGACGGTGTACAGCAATG |
| igtp | CCGTGAACAAGTTCCTCAGGCT | GAGGTCTTGGTGTTCTCAGCCA |
| gfap | GGA GAG GGA CAA CTT TGC AC | CCA GCG ATT CAA CCT TTC TC |
| zbp1 | GATCTACCACTCACGTCAGGAAG | GGCAATGGAGATGTGGCTGTTG |
| 18s | GTA ACC CGT TGA ACC CCA TT | CCA TCC AAT CGG TAG TAG CG |
