## Supplementary figure legend for "IRF3 regulates neuroinflammatory responses and the expression of genes associated with Alzheimer’s disease"

**Supplementary Figure 1. Myeloid infiltration and inflammasome activation could not be detected in the cortical sample of mice injected with a single LPS or 4-day repeated LPS challenge.**

A) Representative images of FACS analysis shows no infiltrating myeloid cells (CD11b^+^,CD45^high^) in the brain in response to 6h of LPS challenge. Arrow points to the microglia gating (CD11b^+^,CD45^intermediate^).

B, C) Western blot images and quantification show no significant increase in pro or cleaved form of IL1β 6h after single LPS (A) (N=4,5) each group or 4 repeated LPS challenges (B) N=9,10 each group. No cleaved isoforms could be detected in any of the conditions.

Two-way ANOVA with Tukey’s multiple comparisons. *p<0.05, **p<0.01, ***p<0.001, ****p<0.0001

**Supplementary Figure 2. 4-day repeated LPS challenge does not compromise BBB integrity in the WT or IRF3 KO mice.**

A) Representative images of western blots of cortical lysates showing expression of BBB proteins- Claudin-1 and Occludin.

B-C) Quantification of the western blots shows no differences in levels of Claudin-1 and Occludin either in the WT or IRF3KO tissue lysates. N=9,10 each group. Two-way ANOVA with Tukey’s multiple comparisons. *p<0.05, **p<0.01, ***p<0.001, ****p<0.0001

**Supplementary Figure 3: Increased IRF3 gene count and the presence of a doublet in IRF3 western blot of EYFP^+^ cells from CNS confirms the expression of IRF3-2D construct.**

A) Quantification of the normalized gene count for IRF3 transcripts from the RNAseq of EYFP+ cells from the brains of mice confirms overexpression of IRF3-2D. IRF3-2D,Cre_Oil mice also showed increased levels of IRF3 transcripts, which were further increased with tamoxifen administration.

N=3,4 each group, *p<0.05, **p<0.01, ***p<0.001. One way ANOVA with, Tukey’s multiple comparisons.

B) Western blot and its quantification from flow-sorted EYFP^+^ cells from IRF3-2D,Cre background shows the presence of a doublet pointed out by arrow indicating the expression of IRF3-2D construct in mice heterozygous for IRF3-2D. Some leaky expression was observed in IRF3-2D,Cre_Oil cells.

**Supplementary Figure 4: Expression of IRF3-2D in microglia does not induce behavioral changes in mice.**

A) Quantification of % time spent in the center or total distance traveled in an open field arena, showed no difference in any of the genotypes tested. N=6-8 per group
B) Quantification of time spent in the open arms was equivalent between all the genotypes tested. N=5-8 per group.
C) Quantification of % alternation in Y-maze test shows no difference in memory performance in any of the genotypes tested. N=6-8 per group

One way Anova, with Sidak’s multiple comparison test.

**Supplementary Figure 5: Leaky IRF3-2D expression leads to inflammatory transcriptional changes.**

A) Volcano plot showing differentially expressed genes in Cre_Oil vs IRF3-2D,Cre_Oil, with a total of 321 differentially expressed genes. (n=2 for Cre_Oil, n=3 for IRF3-2D,Cre_Oil)
B) Gene ontology analysis (biological process) and Reactome analysis of the upregulated genes in A) shows activation of pathways related to leukocyte migration and chemotaxis (yellow arrows), activation of viral response (black arrows antigen presentation (red arrow), immune response pathways (light blue arrow), NOD like receptor signaling pathway (Green arrow) and ER-phagosome pathway (purple arrow).
