## Supplementary figures for "IRF3 regulates neuroinflammatory responses and the expression of genes associated with Alzheimer’s disease"

Supplementary Figure 1: Myeloid infiltration and inflammasome activation could not be detected in the cortical sample of mice injected with a single LPS or 4-day repeated LPS challenge.

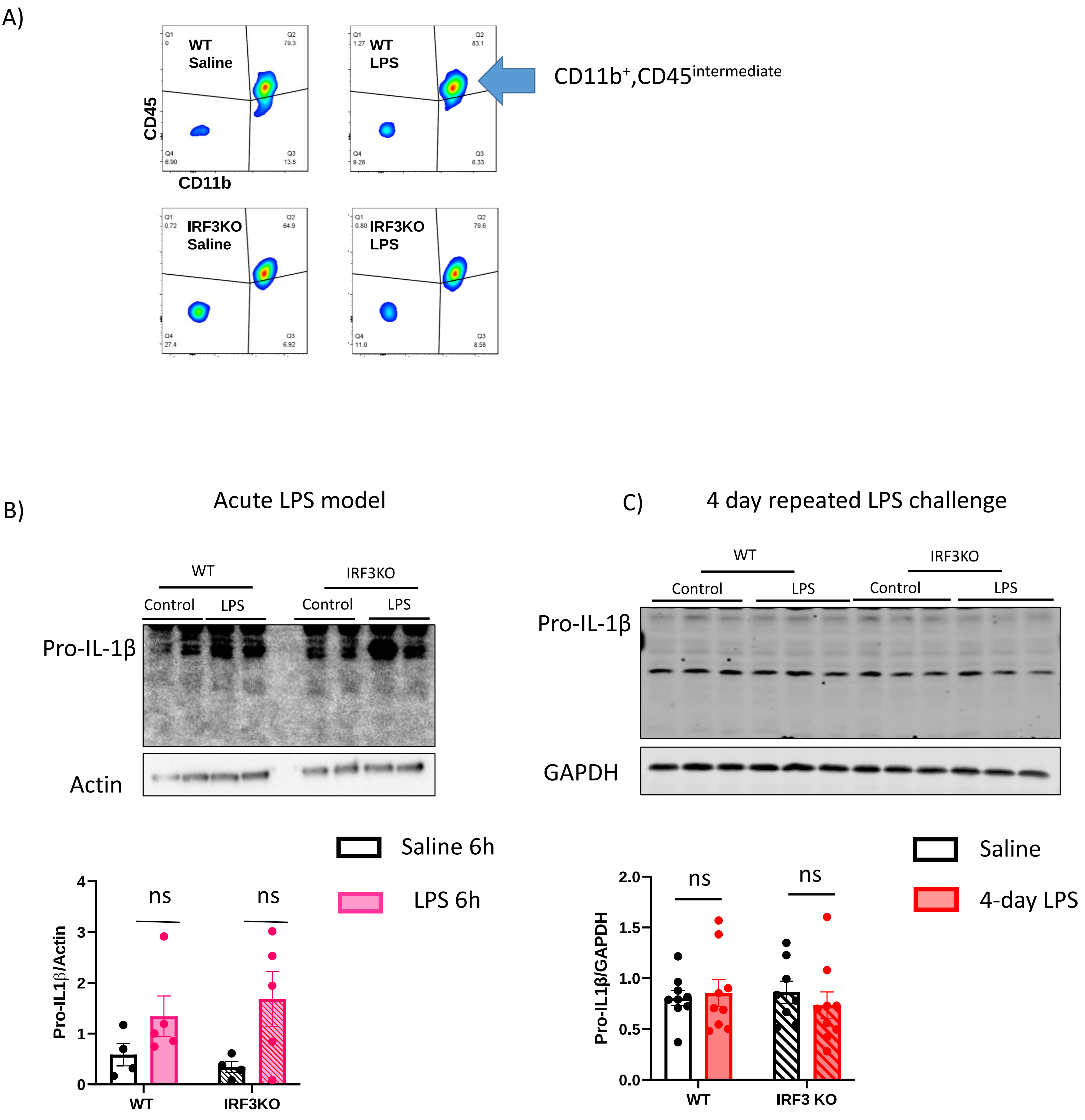

Supplementary Figure 2: 4-day repeated LPS challenge does not compromise the BBB integrity in the WT or IRF3KO mice.

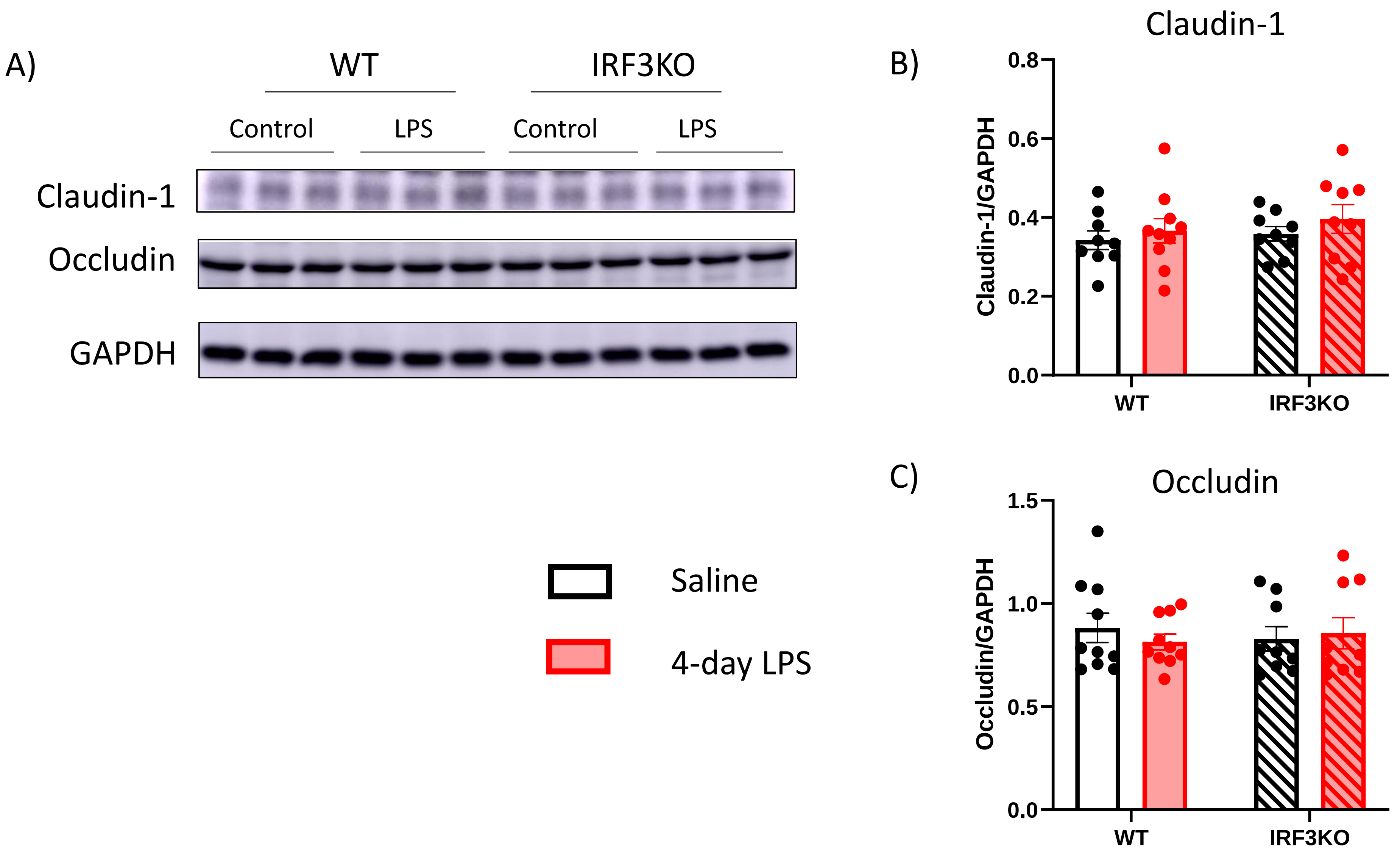

Supplementary Figure 3: Increased IRF3 gene count and the presence of a doublet in IRF3 western blot of EYFP$^{+}$ cells from CNS confirms the expression of IRF3-2D construct.

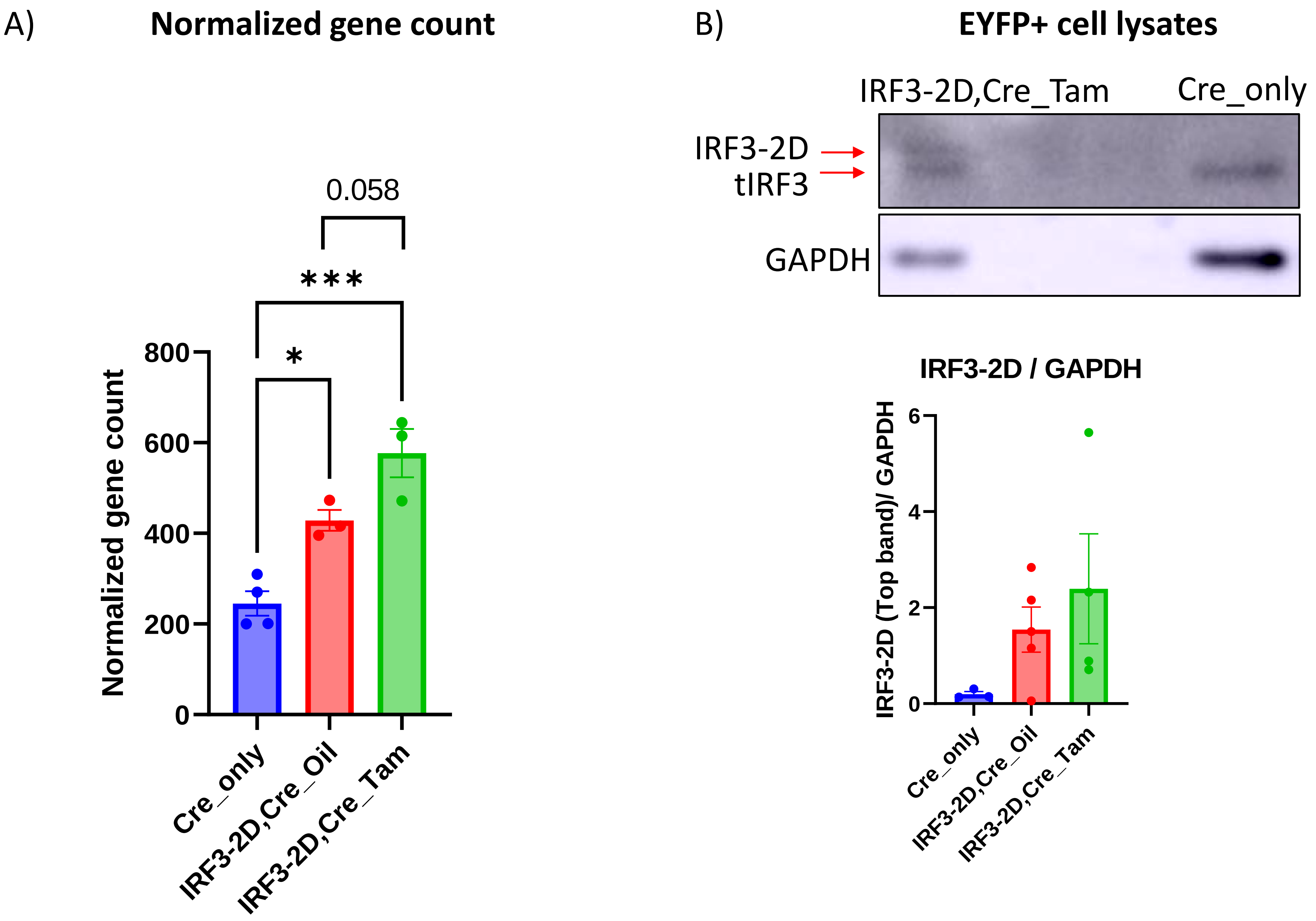

Supplementary Figure 4 :Expression of IRF3-2D in microglia does not induce behavioral changes in mice.

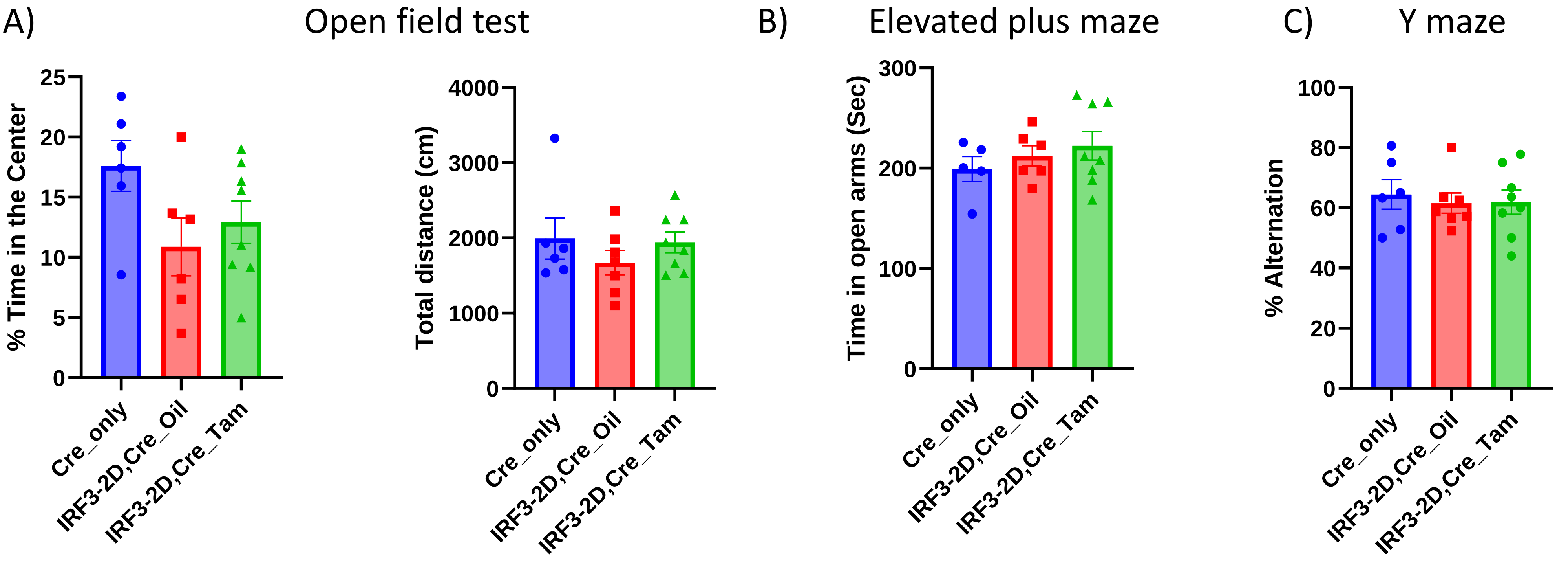

Supplementary Figure 5: Leaky IRF3-2D expression leads to inflammatory transcriptional changes.

A) **Comparison: Cre_Oil vs IRF3-2D,Cre_Oil**

**Volcano plot: q-value** **<u>Differential gene expression count : 321</u>**

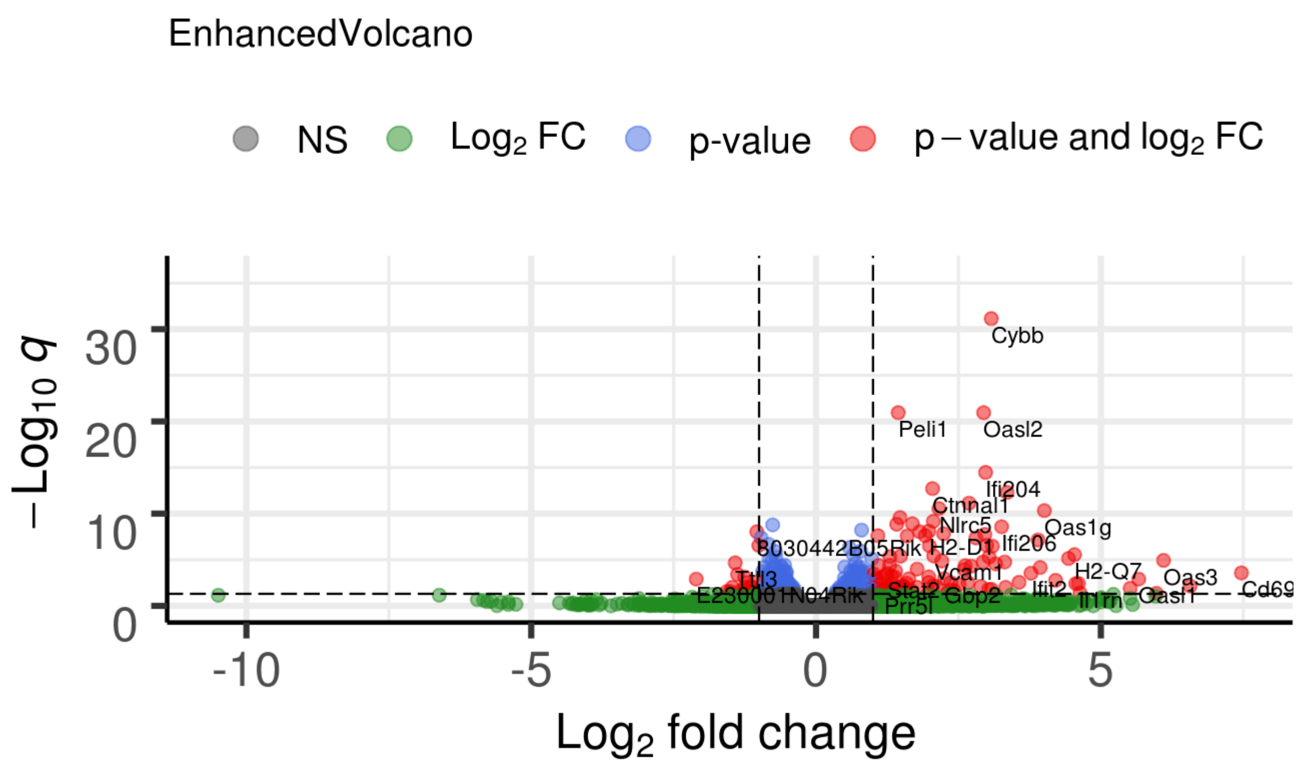


B) **Comparison: Cre_Oil vs IRF3-2D,Cre_Oil**

GO analysis, Biological process

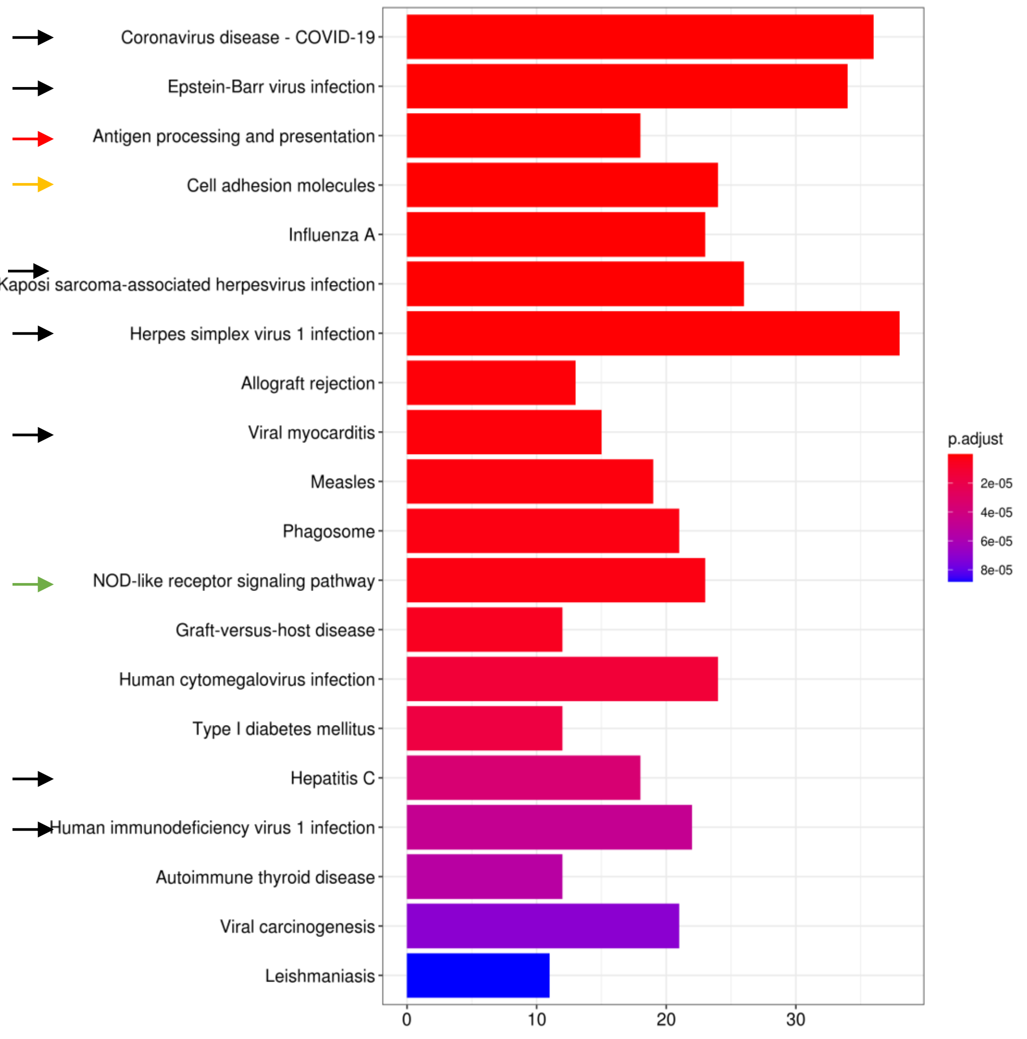


Reactome

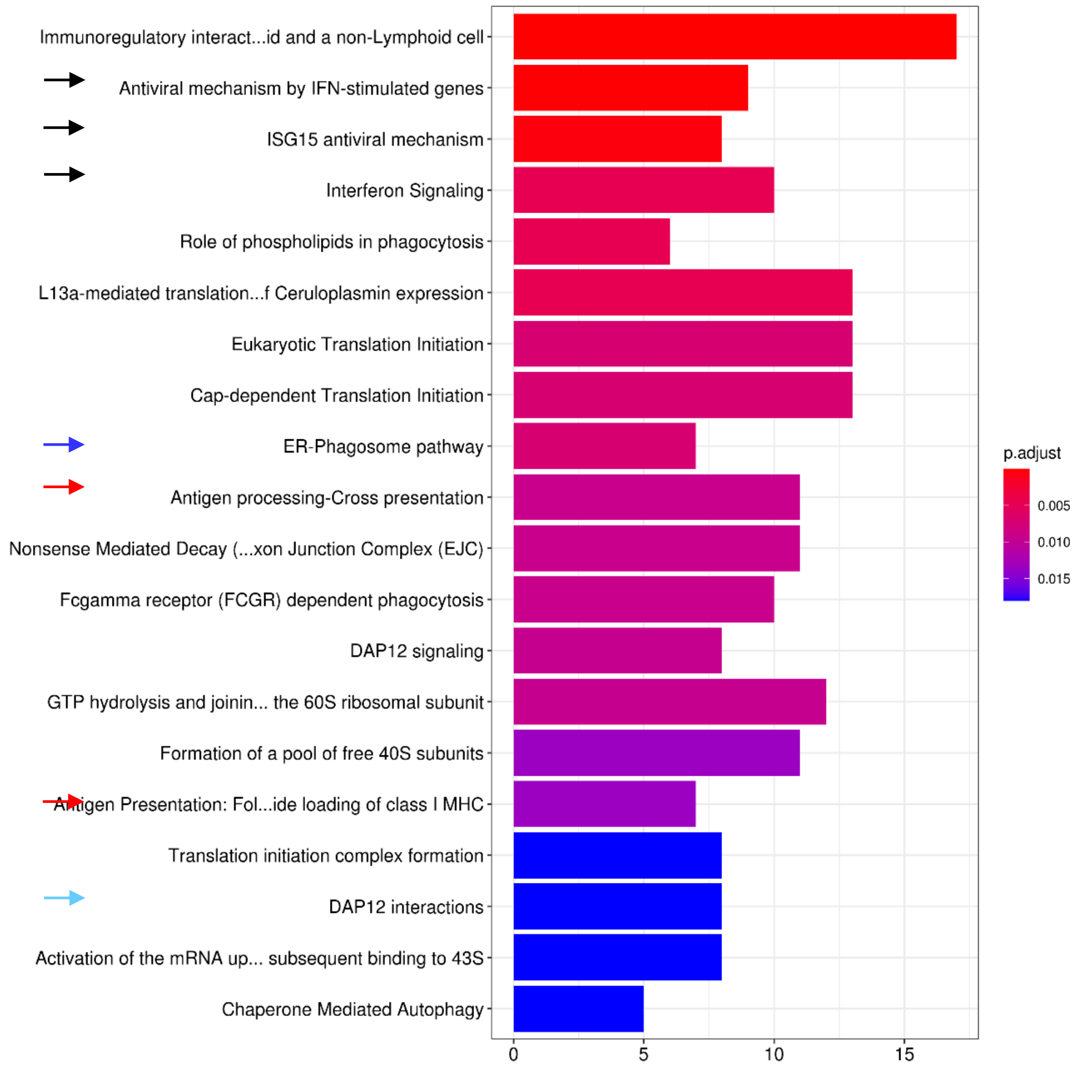
